# The breast tumor microenvironment exploits eosinophil plasticity to suppress their anti-tumor activity

**DOI:** 10.64898/2026.01.12.698961

**Authors:** Zofia Varyova, Mathilde Pohin, Gracie J. Mead, Libby K. Jennings, Valeria da Costa, Sarah Spear, Iain A. McNeish, Anja Schwenzer, Adrian L. Harris, Audrey Gérard, Kim S. Midwood

## Abstract

Eosinophils recently emerged as mediators of anti-tumor immunity in immune checkpoint blockade (ICB) treated breast cancer patients. Yet, their role in the treatment-naïve breast tumor microenvironment (TME) remains elusive. Here, we show that the breast TME shapes eosinophils into a less active state characterized by loss of Ly6C. While bone marrow and circulating eosinophils are Ly6C⁺, this population progressively transitions into a Ly6C⁻ state marked by reduced cytotoxicity and interferon (IFN) responsiveness during tumor progression. Further investigation of Ly6C uncovered previously unappreciated granularity of eosinophil differentiation in vitro, recapitulating the Ly6C⁺ to Ly6C⁻ transition and associated functional loss observed in vivo. IFN stimulation partially restored the Ly6C⁺ phenotype ex vivo. Importantly, in ICB-treated tumors, Ly6C^+^ eosinophils positively correlated with increasing levels of IFNs, suggesting an additional mechanism by which IFNs contribute to effective ICB responses. We propose Ly6C as a key marker of eosinophil differentiation and activation, with the TME shaping eosinophils into a less cytotoxic Ly6C⁻ state.

## Introduction

The interplay between immune and cancer cells within the tumor microenvironment (TME) is a decisive factor in disease progression and therapeutic outcome. While immune checkpoint blockade (ICB) aimed at reactivating cytotoxic T cells has revolutionized treatment for a subset of patients, the mechanisms leading to resistance or to durable response remain only partially understood^1,2^. Eosinophils, granulocytes mostly studied in the context of allergy and parasite infections^3^, recently emerged as important players in a favorable response to ICB in melanoma, non-small cell lung cancer, and triple-negative breast cancer^4–6^. These clinical observations are supported by mechanistic studies demonstrating that eosinophils actively contribute to ICB efficacy in breast cancer models^6,7^. However, while eosinophils exhibit anti-tumorigenic behavior in treatment-naïve colon and lung cancer models^8–11^, their depletion in models of breast cancer had only minor impact on tumor growth unless combined with treatment^6,7,12,13^. This discrepancy raises the question of how the breast TME affects eosinophil function, and which signals enable eosinophil transition from bystanders to active mediators of anti-tumor immunity.

Eosinophils were long thought to emerge from bone marrow as already terminally differentiated granulocytes, with a short half-life upon tissue entry^14^. However, this dogma is challenged by studies showing that eosinophils can be long-lived^15^ and form a plastic population that adapts to their local tissue niche^16^. A recent transcriptomic study revealed a developmental trajectory consisting of precursor and immature eosinophil subsets emerging from the bone marrow, altering their transcriptomic signature in circulation, and then adapting to the gastrointestinal tract by acquiring either basal (matrix modulating) or active (immunomodulatory) phenotypes^17^. Consistent with this plasticity, tumor-associated eosinophils (TAEs) that play an anti-tumorigenic role in colorectal cancer and lung metastasis models display altered transcriptomes compared to their healthy counterparts^9,10,18^. These studies demonstrate that eosinophils are capable of promoting T cell activation within the TME, while also possessing direct cytotoxic potential. The cytotoxic potential mediated by the release of cationic eosinophil granules is well described in responses to helminth infections^16,19^, with recent evidence demonstrating that eosinophils are also capable of killing cancer cells in an adhesion-dependent manner when stimulated with cytokines such as IL-33 or IL-18^20,21^. However, the specific phenotypic trajectories eosinophils undergo within the breast TME, and how these states correlate with their cytotoxic function, remain poorly defined. Therefore, identifying eosinophil phenotypical heterogeneity within the TME is essential for understanding how eosinophils maintain or lose anti-tumor potential and how these states influence their ability to mediate ICB responses.

Here we investigate how eosinophils adapt their phenotype to the progressing TME in murine models of breast cancer. We identified the NT193 tumor model^22^ as unexpectedly eosinophil-rich, which enabled us to mechanistically dissect eosinophil dynamics in wild-type animals without the necessity of using IL-5Tg mice with systemic eosinophilia^23^, previously used to elucidate role of eosinophils in other cancer models^8,10,11,24,25^. We found that bone marrow-resident and circulating eosinophils express relatively high levels of Ly6C, however, upon entering tumors they lose Ly6C expression over time. This tumor specific Ly6C^−^ subset stems from circulating Ly6C^+^ eosinophils and is associated with reduced IFNγ/IFNβ responsiveness and cytotoxicity. Strikingly, Ly6C marked distinct stages of bone marrow-derived eosinophil (BMDEs) development, with virtually all BMDEs losing Ly6C expression under prolonged homeostatic IL-5 signaling, and functionally mirroring loss of IFN responsiveness and cytotoxicity. Therefore, we propose Ly6C as a previously unrecognized eosinophil marker, distinguishing functionally distinct eosinophil subsets. Importantly, IFN signaling enhanced eosinophil cytotoxicity ex vivo, and selectively supported the presence of the Ly6C^+^ eosinophil subset in vivo during ICB treatment, underscoring the potential relevance of the IFN-eosinophil axis for effective ICB response.

## Results

### Late stages of tumor development are characterized by the presence of a Ly6C^−^eosinophil subset

Eosinophils are increasingly recognized as key mediators of anti-tumor immunity across multiple cancer types, acting either through direct cytotoxicity^8,10^ or by mediating T cell responses^9,11^. Yet their role in treatment-naïve breast cancer remains poorly defined, partially due to their low abundance in murine models of this disease. To establish an experimental system suitable for the investigation of eosinophils in breast cancer, we compared eosinophil infiltration of three orthotopic breast cancer models - E0771, 4T1, and NT193 – representing distinct molecular subtypes and host backgrounds. NT193 tumors on the FVB background exhibited a robust eosinophil infiltration, with eosinophils accounting for up to 20% of CD45⁺ immune cells, significantly higher levels than in E0771 or 4T1 tumors (Fig. 1A), with this effect not being driven by an expansion of the myeloid compartment (CD11b^+^) compared to other examined breast cancer models (Supplementary Fig. 1A). To our knowledge, such pronounced eosinophil infiltration was not previously reported in any other wild-type murine breast cancer model. Therefore, the NT193 model provides a unique opportunity to dissect eosinophil biology in the breast TME.

**Figure 1.**
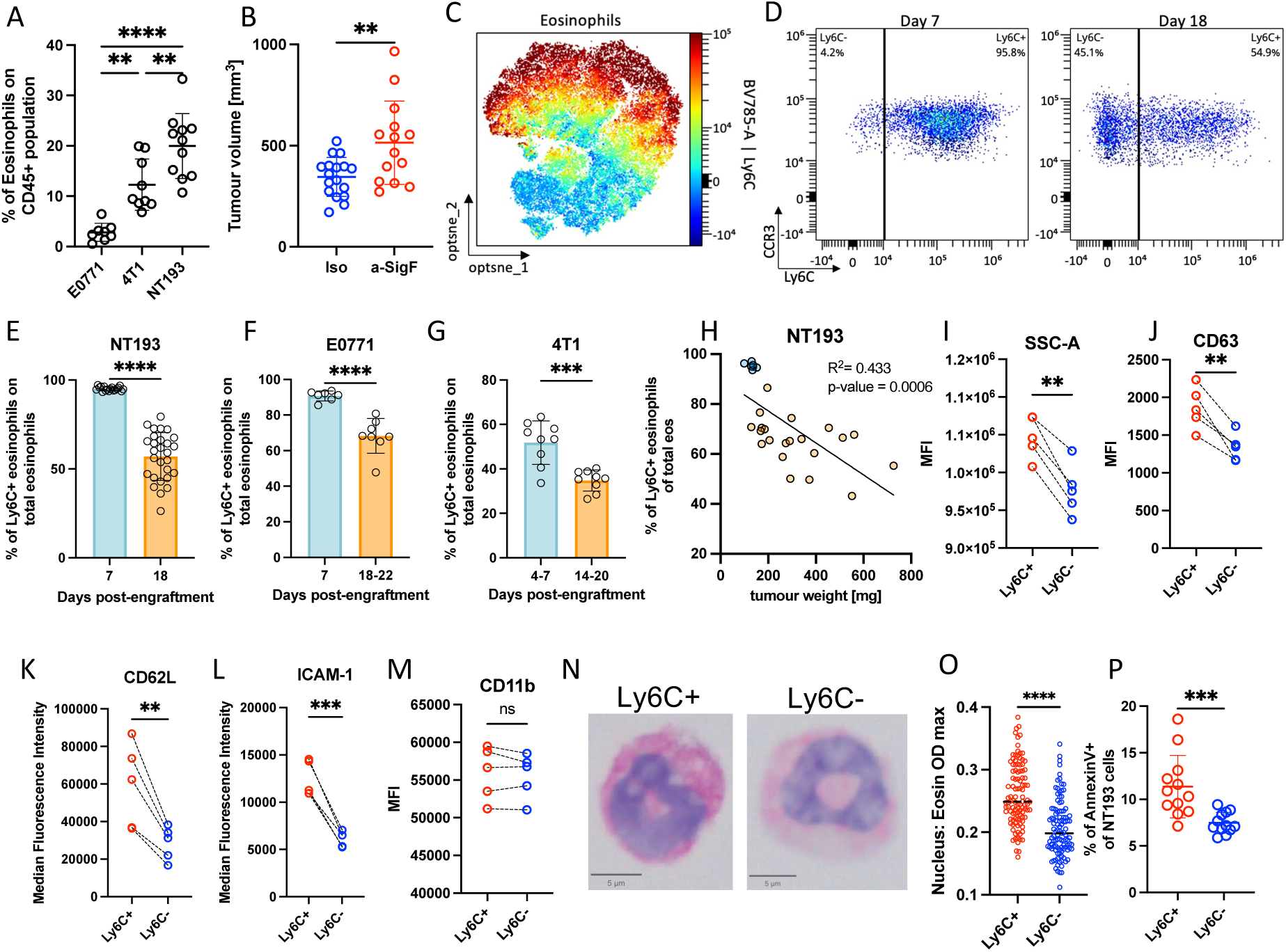
Tumor progression leads eosinophils into a less cytotoxic Ly6C- state. **(A)** Infiltration of eosinophils at the final time point in the NT193 (day 18, n=11), 4T1 (day 17-20, n=9), and E0771 (day 18-22, n=8) orthotopic mammary tumors engrafted into the 4^th^ mammary fat pad of FVB, Balb/c, and C57/BL6 females, respectively. Data pooled from 2 independent experiments. **(B)** Tumor volume of the NT193 tumor-bearing mice that were treated with an isotype control (n = 16) or an anti-Siglec-F (n=15). Data pooled from 3 experiments. **(C)** opt-SNE heatmap of Ly6C eosinophil expression analyzed in NT193 tumors on day 18. **(D)** Representative flow cytometry plots of tumor-infiltrating eosinophils (Siglec-F^+^, Ly6G^−^) from NT193 tumors harvested on day 7 and day 18. Data representative of at least 3 independent experiments. **(E-G)** Proportion of Ly6C^+^ eosinophils of total eosinophil population in NT193 (n_day7_=7, n_day18_=18), E0771 (n_day7_=7, n_day18_=8), 4T1 (n_day4-7_=9, n_day14-20_=9) orthotopic breast cancer models on indicated days, analyzed by flow cytometry. Data pooled from 2 independent experiments per condition. **(H)** Correlation between Ly6C^+^ eosinophils and tumor weight of NT193 tumors harvested on day 7 (blue, n=7) and day 18 (orange, n=20). Data pooled from 2 independent experiments. **(I-M)** Comparison of median fluorescence intensity of SSC-A **(I)**, CD63 **(J)**, CD62L **(K)**, ICAM-1 **(L)**, and CD11b **(M)** between matched Ly6C^+^ and Ly6C^−^ eosinophils from NT193 tumors on day 18 (n=5). Data representative of 2 independent experiments. **(N)** Representative images of cytospun Ly6C^+^ and Ly6C^−^ eosinophils sorted from NT193 tumors on day 18 stained with H&E. **(O)** Quantification of eosin optical density of cytospun eosinophils (n_Ly6C+_ = 106, n_Ly6C-_ = 107). Data representative of 5 out of 6 tumors. **(P)** Proportion of apoptotic Annexin-V+ NT193 cells after direct co-culture with Ly6C^+^ or Ly6C^−^ eosinophils sorted from NT193 tumors on day 18. Data are pooled from 3 independent experiments (n_Ly6C+_ = 12, n_Ly6C-_ = 12). Data show individual values and mean or mean ± SD, and were analysed by unpaired Student’s t-test, or by 2-way ANOVA for comparison of two and more groups. Statistical significance is displayed on figures as follows: *p < 0.05, **p < 0.01, ***p<0.001, ****p<0.0001.

To determine the role of eosinophils in the NT193 tumors, tumor-bearing mice were treated with an anti-Siglec-F antibody that efficiently depleted circulating and tumor-resident eosinophils (Supplementary Fig. 1B, C). Eosinophil depletion resulted in a modest but significant increase in the tumor volume (Fig. 1B), consistent with the anti-tumorigenic potential of these cells described in other cancer models^8–12^. The difference in tumor growth became more pronounced at the later time points (Supplementary Fig. 1D), coinciding with increased eosinophil relocation from the periphery to the tumor core (Supplementary Fig. 1E, F). Together, these data show that eosinophils can restrain late-stage tumor growth.

To explore if eosinophils display phenotypic heterogeneity during tumor progression, we performed high-dimensional flow cytometry analysis of early and late NT193 tumors, using a set of lineage markers of myeloid populations (Siglec-F, CD11b, F4/80, Ly6G, Ly6C), eosinophil chemokine/alarmin receptors (CCR3, IL33r, CCR5), maturation and activation markers (IL5Ra, CD11c) and the antigen-presenting marker (MHC-II) (Supplementary table 1). This analysis identified two phenotypically distinct eosinophil subsets distinguished by Ly6C expression (Fig. 1C). Both Ly6C^+^ and Ly6C^−^ subsets expressed similar levels of the canonical maturity and activation markers examined (Supplementary Fig. 1G). Interestingly, Ly6C^−^ eosinophils emerged exclusively during the late stages of NT193 tumor development (Fig. 1D, E). The presence of the Ly6C^+^ subset was further validated in early and late stage E0771 and 4T1 tumors (Fig. 1F, G; Supplementary Fig. 1H, I), indicating that the appearance of Ly6C^−^ eosinophils is a well-conserved feature of the late TME. Moreover, the temporal distribution of these two populations was conserved in an ovarian cancer model ID8^26^, unrelated to mammary tissue, with the Ly6C^−^ eosinophil subset being exclusively associated with tumor progression (Supplementary Fig. 1J). The association of Ly6C downregulation with tumor progression across various cancer models, and the negative correlation of the proportion of Ly6C^+^ eosinophils with tumor volume in NT193 tumors (Fig. 1H), suggested that the Ly6C^+^ subset might be responsible for eosinophil-mediated anti-tumor activity.

We therefore further investigated the activation state of these two subsets more in depth. Ly6C^+^ eosinophils displayed higher expression of markers associated with increased cytotoxicity^11,17,21^, including increased granularity, elevated expression of the degranulation marker CD63, and adhesion markers selectin CD62L and ICAM-1, markers potentially enabling direct interactions of activated eosinophils with tumor cells, while CD11b levels were comparable to Ly6C^−^ eosinophils (Fig. 1I-M). Despite the phenotypic differences, both subsets exhibited similar nuclear morphology (Fig. 1N), suggesting that the observed heterogeneity stems from distinct activation states rather than maturation differences. Consistent with the increased granularity observed by flow cytometry, Ly6C^+^ eosinophils displayed a higher intensity of eosin staining, while the hematoxylin intensity remained unchanged (Fig. 1O, Supplementary Fig. 1K). To confirm the increased cytotoxic capacity of Ly6C^+^ eosinophils, the cytotoxic potential of tumor-associated eosinophils (TAEs) sorted from NT193 tumors was tested by direct co-culture with the NT193 cell line. Ly6C^+^ eosinophils induced tumor cell apoptosis more efficiently compared with the Ly6C^−^ subset (Fig. 1P), further establishing their higher anti-tumorigenic potential.

Together, these results demonstrate phenotypical heterogeneity of TAEs that is affected by the progressing TME, with Ly6C expression marking a more cytotoxic subset whose frequency declines as tumors progress, suggesting that the TME drives eosinophils toward a less cytotoxic, Ly6C^−^ state.

### Ly6C^+^ eosinophils transition into the Ly6C^−^ subset in conditions supporting their survival

To track the origin of Ly6C^−^ eosinophils associated with late-stage tumor development, we investigated the expression of Ly6C on eosinophils in the bone marrow, blood, and mammary fat pad of healthy and tumor-bearing mice. To assess the relative Ly6C expression on eosinophils, we directly compared them with monocytes (Ly6C^high^, SiglecF^−^, Ly6G^−^) and neutrophils (Ly6C^+^, SiglecF^−^, Ly6G^+^), cell types with well-characterized Ly6C expression^27,28^, across the selected tissues (Supplementary Fig. 2A-D). Furthermore, because eosinophils are bone marrow-derived granulocytes that infiltrate mammary tissues along the CCL11-CCR3 axis^29^, CCR3 was used as a marker of eosinophil maturation and migration.

In the bone marrow, eosinophils uniformly expressed high levels of Ly6C both in the presence or absence of tumors, and were further divided into immature (CCR3^−^) and mature (CCR3^+^) populations (Fig. 2A). As eosinophils emerged from the bone marrow, only the mature CCR3^+^ Ly6C^+^ population was detected in the circulation regardless of tumor status, indicating that Ly6C downregulation does not occur in circulation. To exclude the possibility that Ly6C^−^TAEs arise from a population of resident eosinophils in mammary tissue, we next examined Ly6C expression on eosinophils in the mammary fat pads of healthy mice (Fig. 2A). Although mammary tissue-resident eosinophils presented with Ly6C^+^ phenotype, they displayed a modest reduction in Ly6C expression compared to neutrophils together with CCR3 downregulation, indicative of internalization as a part of adaptation to the tissue following eotaxin mediated chemotaxis^30–32^. However, given that this resident population accounted for less than 1% of all viable mammary resident cells, and only around 5% of the immune compartment within the healthy fat pad, this population is unlikely to significantly contribute to either of the TAE subsets accumulating in the tumors throughout disease progression (Supplementary Fig. 3A, B). Finally, as shown above (Fig. 1D), TAE downregulate Ly6C expression during tumor progression, with both Ly6C^+^ and Ly6C^−^ subsets being CCR3^−^, suggestive of their tissue resident phenotype (Fig. 2A). These findings collectively indicate that Ly6C downregulation does not reflect cancer-induced reprogramming of eosinophil development in the bone marrow, or in the circulation, but rather that this change arises within the TME following tissue infiltration.

**Figure 2.**
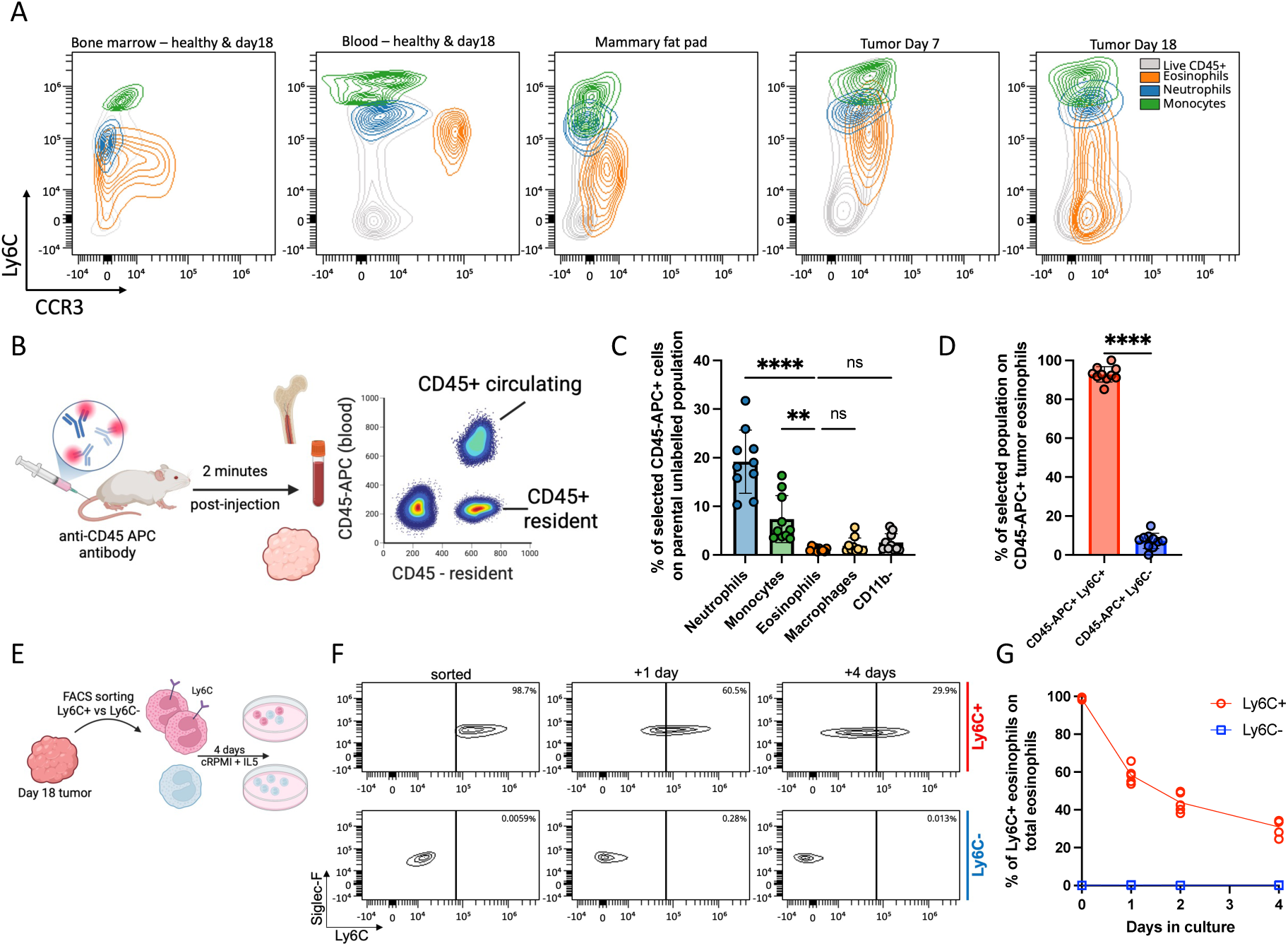
Ly6C^+^ eosinophils transition to the Ly6C^−^ subset in situ and ex vivo. **(A)** Representative flow cytometry contour plots comparing CCR3 and Ly6C expression on the total population of lymphocytes (grey), eosinophils (orange), neutrophils (blue), and monocytes (green) in mammary fat pads of healthy FVB mice, and bone marrow, blood, and tumors of the NT193 tumor-bearing mice on the indicated days. Representative of 3 independent experiments. **(B)** Schematics of in vivo labelling by anti-CD45-APC antibody and the flow cytometry analysis of the harvested bone marrow, blood and tumours of CD45-APC labelled animals. Graphics created with BioRender. **(C)** Proportion of circulating CD45-APC+ tumor-infiltrating neutrophils, monocytes, eosinophils, macrophages, and non-myeloid (CD11b-) cells (n = 10). Data pooled from 2 independent experiments, analyzed by one-way ANOVA comparisons to eosinophil population. **(D)** Proportion of CD45-APC labelled Ly6C^+^ and Ly6C^−^eosinophils of total CD45-APC^+^ eosinophil population in the NT193 tumors between day 15-18, (n = 10). Data pooled from 2 independent experiments. **(E)** Schematic representation of experimental design, Ly6C^+^ and Ly6C^−^ eosinophils were sorted from the NT193 tumors on day 18 and cultured up to 4 days in the presence of IL-5. Graphics created with BioRender. **(F)** Representative contour plots of Ly6C expression of sorted Ly6C^+^ and Ly6C^−^ eosinophils on the indicated time points. **(G)** Percentage of the Ly6C^+^ eosinophils in ex vivo cultures of sorted eosinophils (n_Ly6C+_ = 5, n_Ly6C-_ = 5). Representative of 3 independent experiments. All data show individual values and mean or mean ± SD, and were analysed by unpaired Student’s t-test, or by 2-way ANOVA for comparison of two and more groups. Statistical significance is displayed on figures as follows: ns > 0.05, ****p<0.0001.

Given that all circulating eosinophils retain Ly6C expression, we hypothesized that the Ly6C^+^ phenotype of TAE may reflect a vascular or recently recruited population, whereas the Ly6C^−^ eosinophils would represent a tumor resident subset. Therefore, we employed intravenous labelling with an anti-CD45-APC conjugated antibody, that enabled us to track circulating eosinophils in bone marrow, blood and tumors of the same animal, a method that allowed us to compare Ly6C expression of the tissue resident and circulating eosinophils of tumor-bearing mice on day 18 (Fig. 3B). Consistent with the proposed mature phenotype of CCR3^+^ eosinophils^33,34^, a subset of CCR3^+^ Ly6C^+^ bone marrow eosinophils were labelled with anti-CD45-APC (Supplementary Fig. 3C), indicating they were in the process of emerging into the circulation at the time of the anti-CD45-APC intravenous labelling. In the blood, virtually all eosinophils were CD45-APC^+^ Ly6C^+^, confirming the effectiveness of the labeling strategy (Supplementary Fig. 3D). In contrast, CD45-APC^+^ eosinophils represented only a minor population of all TAE (<2%) (Fig. 2C, Supplementary Fig. 3D), compared to other myeloid cells more abundant in the tumor vasculature, such as tumor-associated neutrophils or monocytes (20% and 8%, respectively). However, within the relatively small circulating subset of TAEs, more than 90% of cells were Ly6C^+^ eosinophils (Fig. 2D), demonstrating that this population is emerging from the vasculature, and consistent with the hypothesis that eosinophils lose Ly6C expression after entering tumors. Moreover, immunofluorescent staining revealed extravascular localization of eosinophils at early and late stage tumor development, consistent with eosinophil tissue residency (Supplementary Figs. 1E, 3E).

**Figure 3.**
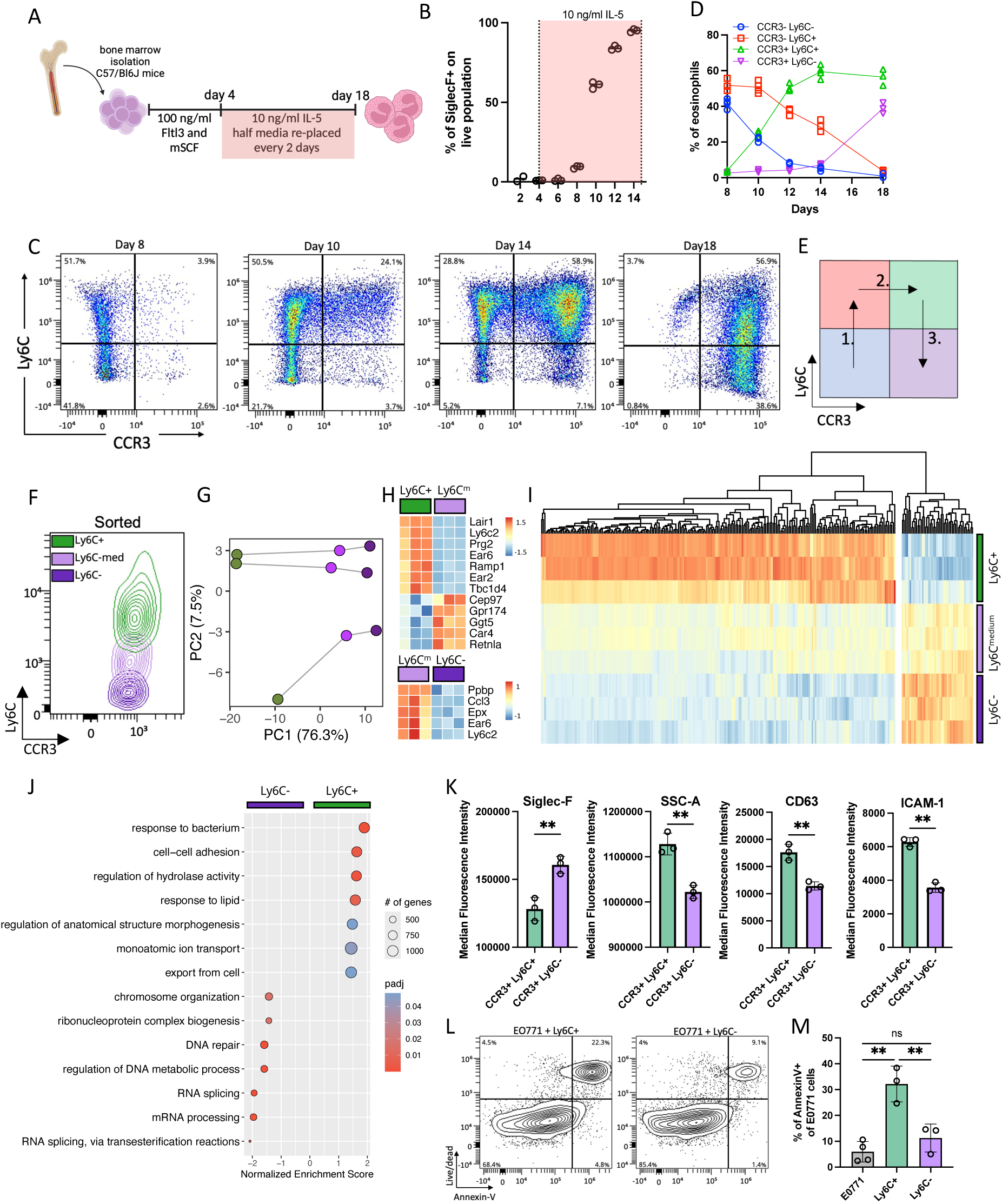
Bone marrow-derived eosinophils recapitulate the Ly6C^+^ transition into the Ly6C^−^subset. **(A)** Schematics of experimental design. Bone marrow progenitors of C57/Bl6 mice were cultured for 4 days in the presence of Fltl3 and mSCF, afterwards, cells were cultured in the presence of IL5. Graphics created with BioRender. **(B)** Percentage of eosinophils of total cells in culture on the indicated days. **(C)** Representative flow cytometry plots of CCR3 and Ly6C expression of bone marrow-derived eosinophils (BMDEs) on the indicated days. **(D)** Time-course analysis of eosinophil development. Percentage of individual eosinophil subsets of the total eosinophil population on the indicated timepoints (n = 3), representative of 2 independent experiments. **(E)** Schematics of proposed eosinophil development in ex vivo bone marrow-derived culture**. (F)** Representative contour plot of sorted BMDE CCR3+ eosinophils based on their Ly6C positivity on day 18 (n = 3). **(G-J)** FACS-sorted BMDEs were analyzed by bulk-RNA sequencing. **(G)** PCA plot of PC1 and PC2 components, of Ly6C^+^, Ly6C^med^, and Ly6C^−^ sorted eosinophils analyzed by bulkRNA-seq on day 18**. (H)** Heatmaps of the top DEGs. **(I)** Heatmap analysis of all DEGs z-scored across all samples**. (J)** GSEA-GO comparison of Ly6C^+^ vs Ly6C^−^ BMDEs. Differences in gene expressions are shown as normalized enrichment score (NES), and signatures are organized by increasing absolute NES values. The size of the circle represents the number of genes identified in the pathway; color represents the adjusted p-value. **(K)** Comparison of median fluorescence intensity of Siglec-F, SSC-A, CD63, ICAM-1 between BMDE Ly6C^+^ and Ly6C- eosinophils on day 18 (n = 3). Data representative of 2 independent experiments. **(L)** Representative flow cytometry plots of the E0771 cell line stained by Annexin-V and fixable viability dye after co-culture with Ly6C^+^ or Ly6C^−^ BMDEs. **(M)** Percentage of Annexin-V+ E0771 cells after direct co-culture with sorted Ly6C^+^ (n = 3) and Ly6C^−^ (n = 3) bone marrow-derived eosinophils. Data representative of 3 independent experiments. Data show individual values + median and were analysed by 2-way ANOVA for comparison of 2 and more groups. Otherwise, data show individual values + median and were analysed by Student’s t-test, or by 2-way ANOVA for comparison of two and more groups. Statistical significance is displayed on figures as follows: *p < 0.05, **p < 0.01, ***p<0.001, ****p<0.0001.

Because eosinophils accumulate in late-stage tumors (Supplementary Fig. 3A, B), we hypothesized that Ly6C^+^ eosinophils represent a population that progressively transitions to the Ly6C^−^ state in an environment that favors the eosinophil survival. To test this, Ly6C^+^ and Ly6C^−^ eosinophils were isolated by fluorescence-activated cell sorting (FACS) from the NT193 tumors and cultured in the presence of IL-5 (Fig. 2E), a cytokine essential for eosinophil survival and expansion^35^. Consistent with this model, Ly6C^+^ eosinophils downregulated Ly6C over time, while Ly6C^−^ eosinophils did not gain Ly6C expression (Fig. 2F, G). Similarly, Ly6C^+^ eosinophils isolated from early tumors displayed the same capacity to transition towards the Ly6C^−^ state (Supplementary Fig. 3F-H).

Overall, these results identify Ly6C^+^ eosinophils as a root cluster that can give rise to the Ly6C^−^ subset in the presence of the pro-survival cytokine IL-5 ex vivo, or within the tissue, such as under the influence of the TME.

### Ex vivo eosinophil development is characterized by differential expression of Ly6C and CCR3

Because Ly6C^+^ TAEs progressively lost Ly6C expression in the breast TME and when cultured ex vivo in the presence of IL-5, we next asked whether Ly6C might be a novel marker reflecting eosinophil maturation/differentiation state. Therefore, we employed a bone marrow-derived eosinophil (BMDE) culture model that recapitulates IL-5-driven eosinophil maturation ex vivo (Fig. 3A). As previously reported^36^, Siglec-F^+^ eosinophils constituted nearly 100% of viable cells from day 14 onwards (Fig. 3B), confirming efficient lineage commitment and differentiation. Flow cytometry analysis of maturing BMDEs over time, starting from the day of the first identifiable eosinophil population, day 8, revealed a continuous trajectory of eosinophil maturation clearly defined by Ly6C and CCR3 expression (Fig. 3C). Ly6C expression distinguished two immature subsets, CCR3^−^ Ly6C^−^ and CCR3^−^ Ly6C^+^ at day 8. Following maturation and acquisition of CCR3, eosinophils maintained Ly6C expression on day 10 (CCR3^+^ Ly6C^+^ subset), before starting to downregulate Ly6C, ultimately resulting in a CCR3^+^ Ly6C^−^ population that is increasingly prevalent by day 18 (Fig. 3D). These transitions occurred in directional manner; sorted CCR3^−^ Ly6C^+^ eosinophils rapidly upregulated CCR3 within 2 days of culture, while sorted CCR3^+^ Ly6C^+^ cells transitioned only to a CCR3^+^ Ly6C^−^ state during culture, and sorted CCR3^+^ Ly6C^−^ eosinophils remained phenotypically stable (Fig. 3E, Supplementary Fig. 4A, B). Consistent with these findings, virtually all BMDEs adopted a CCR3^+^ Ly6C^−^ phenotype by day 30 of ex vivo culture, coinciding with declining viability (Supplementary Fig. 4C,D). Together these data indicate that the CCR3^+^ Ly6C^−^ population is a terminally differentiated eosinophil subset.

To gain insights into the differences between these transitioning populations, and to assess the impact of the loss of Ly6C on eosinophil phenotype during development, we sorted mature CCR3^+^ BMDEs into 3 populations: Ly6C^+^, Ly6C^med^, and Ly6C^−^ subsets (Fig. 3F), and performed bulk RNA sequencing (bulkRNA-seq). Principal component analysis revealed a continuous transcriptional trajectory from Ly6C^+^ through Ly6C^med^ to the Ly6C^−^ state (Fig. 3G), consistent with the phenotypic progression observed by flow cytometry. Differential expression analysis confirmed significant transcriptional differences between Ly6C^+^ BMDEs and both Ly6C^med^ and Ly6C^−^ BMDEs, while Ly6C^med^ and Ly6C^−^ eosinophils displayed relatively high similarity (Supplementary Fig. 4E). Examination of the top differentially expressed genes (DEGs) validated the sorting strategy, as *Ly6c2* expression gradually decreased from Ly6C^+^ to Ly6C^−^ BMDEs (Fig. 3H, Supplementary Table 2). Moreover, while Ly6C^+^ BMDEs expressed higher levels of eosinophil granule proteins such as *Prg2 and Ear2/6*, the Ly6C^−^ subset presented with higher expression of tissue repair molecules such as *Retnla*. Global DEG analysis further supported the directional transition, showing a smooth continuum from Ly6C^+^ to Ly6C^−^ eosinophils, with Ly6C^med^ cells representing an intermediate, transcriptionally indistinct state from the Ly6C^−^ subset (Fig. 3I, Supplementary Fig. 4E). On this basis, we focused the subsequent analysis on comparison between Ly6C^+^ and Ly6C^−^ BMDE subsets.

Gene set enrichment analysis using the GO resource (GSEA-GO) (Supplementary Table 3) pointed at Ly6C^+^ eosinophils upregulating pathways linked to “*response to bacterium*” driven by strong overexpression of eosinophil cationic proteins and pro-inflammatory cytokines (*Epx*, *Prg2*, *Ear2*, *Cxcl2, Cxcl3, Il1a), as well as* pathways associated with “*cell-cell adhesion*” related to expression of adhesion molecules (*Icam1*, *Sell* (CD62L), *Cd92*) (Fig. 3J). These results are consistent with the more activated and adhesive phenotype described on the protein level for Ly6C^+^ TAE (Fig. 1I-L). In contrast, Ly6C^−^ eosinophils presented with enrichment of pathways associated with “*DNA repair*” and “*mRNA processing*”. Multiple canonical genes associated with senescence (*Cdkn1a*, *Pml*, *Atm*, *Atr*, *Chek2*, *Prkdc*) were consistently upregulated in the Ly6C^−^ subset (Supplementary Fig. 4F). Although not reaching statistical significance, the GSEA suggested a modest bias towards cellular senescence in Ly6C^−^ BMDEs (Supplementary Fig. 4G). Together, these transcriptional signatures indicate that the pathways elevated in Ly6C- BMDEs may reflect features of senescence associated with the proposed terminal differentiation.

The enrichment of genes encoding for eosinophil granule proteins and activation molecules in Ly6C^+^ BMDEs prompted us to further validate functional markers across these subsets. Consistent with the maturation continuum, Siglec-F expression increased with Ly6C loss. Moreover, granularity, CD63 and ICAM-1 expression were significantly increased in the Ly6C^+^ state, indicating an elevated cytotoxic and adhesive potential (Fig. 3K). In line with these data, direct co-culture of Ly6C^+^ and Ly6C^−^ BMDEs with a strain-matched E0771 breast cancer cell line revealed that only Ly6C^+^ eosinophils were capable of inducing tumor cell apoptosis above baseline levels (Fig. 3L, M).

Thus, our data provide novel evidence that ex vivo differentiation of bone marrow progenitors under IL-5 homeostatic signaling provides a platform to dissect eosinophil maturation in greater detail than previously thought. Importantly, BMDEs recapitulated the Ly6C^+^ to Ly6C^−^ transition observed in TAEs, the key phenotypical differences between these two eosinophil subsets, and further confirmed the link between Ly6C downregulation and loss of cytotoxic potential in an independent model. Collectively, these findings further establish Ly6C as a previously unrecognized marker of eosinophil maturation/differentiation.

### Ly6C^+^ eosinophils represent an IFN-responsive subset, with IFNs enhancing eosinophil cytotoxicity

Data generated up to this point reveal Ly6C expression as a marker of eosinophil differentiation trajectory that is consistently observed in situ, ex vivo, and within in vitro models, in which it coincides with loss of cytotoxicity. However, how the TME further functionally programs eosinophils beyond this intrinsic differentiation process remained unclear. Therefore, we next profiled FACS-sorted Ly6C^+^ and Ly6C^−^ TAE by bulkRNA-seq (Fig. 4A). PCA analysis confirmed subset differences on the transcriptional level (Supplementary Fig. 5A), and subsequent differential expression analysis revealed 116 DEGs (log2FC > 1, padj < 0.05) (Supplementary Table 4), with *Ly6c2* among the most upregulated genes in Ly6C^+^ subset (Supplementary Fig. 5B). GSEA-GO analysis highlighted upregulation of gene sets such as “*killing of cells of another organism*” in Ly6C^+^ eosinophils, consistent with their anti-tumorigenic potential ex vivo. Moreover, the most differentially upregulated pathways in the Ly6C^+^ eosinophil subset were associated with “*response to type II interferon* (IFNγ)” and “*response to interferon beta* (IFNβ)” (Fig. 4B, Supplementary Table 5). The previously reported increase of IFN responsiveness of TAE across multiple studies^9,10,18^, the link between IFNγ stimulation and increased eosinophil cytotoxicity against colorectal cancer cell lines^10^, and the STAT1-dependent Ly6C expression upon IFN stimulation^37^, prompted us to investigate the potential effect of IFNs on Ly6C^+^ and Ly6C^−^ eosinophil behavior.

**Figure 4.**
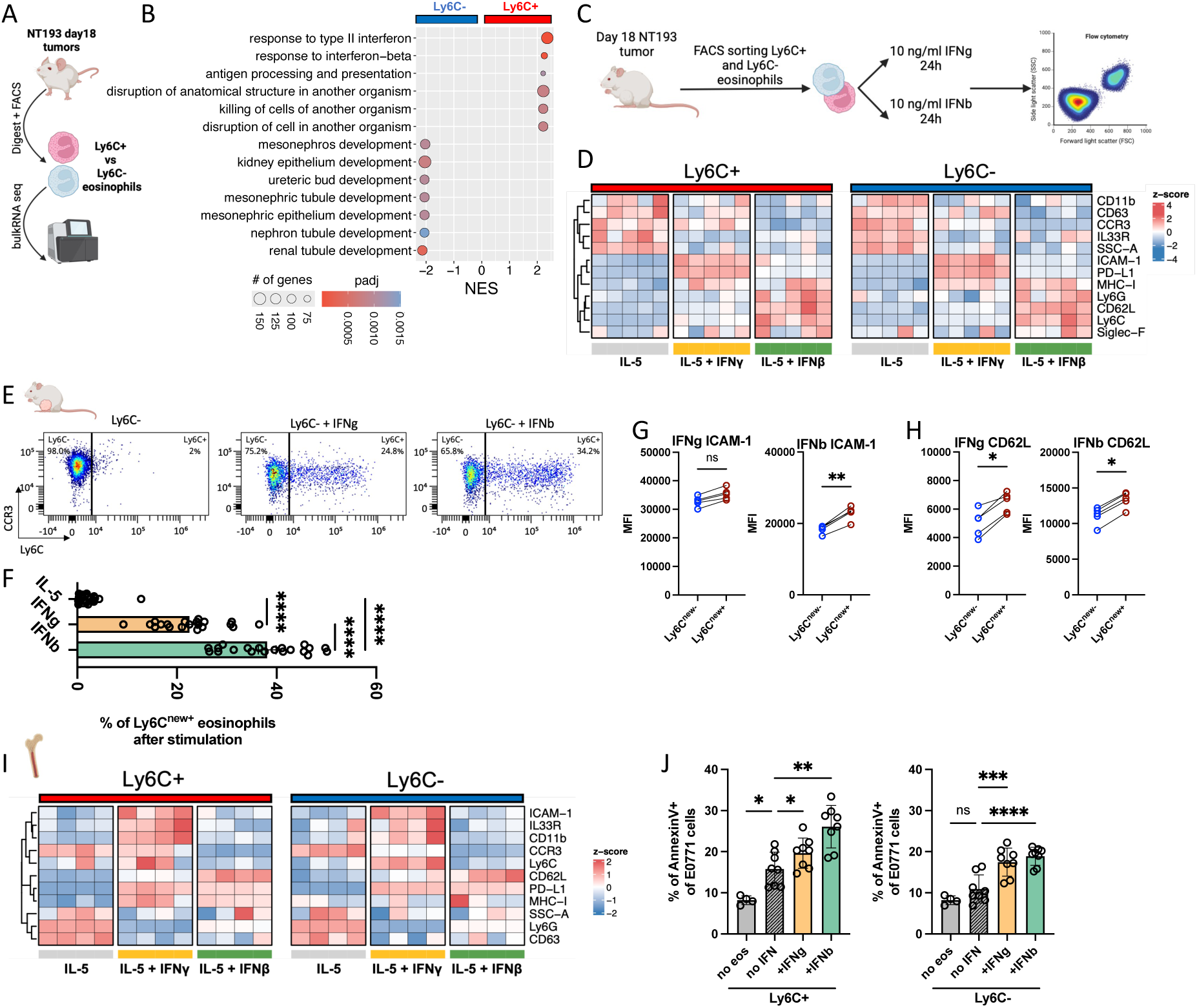
Ly6C^−^ eosinophils lose IFN responsiveness, with IFNs affecting eosinophil activation and cytotoxic potential. **(A)** Schematics of the experimental plan. The Ly6C^+^ and Ly6C^−^ eosinophils were FACS-sorted from the NT193 tumors and analyzed by bulkRNA-seq. Graphics created with BioRender. **(B)** GSEA-GO signatures upregulated in Ly6C^+^ and Ly6C^−^ eosinophils. Differences in gene expressions are shown as normalized enrichment score (NES), and signatures are organized by increasing absolute NES values. The size of the circle represents the number of genes identified in the pathway; color represents the adjusted p-value. **(C)** Schematics of experimental approach. Ly6C^+^ and Ly6C^−^ BMDE and tumor-associated eosinophils stimulated by IFNγ or IFNβ were analyzed by flow cytometry. Graphics created with BioRender. **(D)** Heatmaps showing z-scored surface marker expression in Ly6C⁺ and Ly6C⁻ TAEs (n = 5) stimulated with IL-5, IL-5+IFNγ, or IL-5+IFNβ. Z-scoring was performed separately for Ly6C⁺ and Ly6C⁻ subsets to highlight subset relative changes upon stimulation. Bottom annotation indicates stimulation conditions (gray = IL-5, yellow = IFNγ, green = IFNβ). **(E)** Representative density plots showing Ly6C expression of the tumor sorted Ly6C^−^ eosinophils following IFN stimulation, as indicated. **(F)** Proportions of Ly6C^new+^ subsets of all viable TAEs after stimulation. Data pooled from 4 independent experiments (n = 20). **(G, H)** Ly6C^−^ eosinophils stimulated with IFNγ or IFNβ were re-gated as indicated in Fig. 4E into Ly6C^new−^ and Ly6C^new+^ subsets, and CD62-L **(G)**, and ICAM-1 **(H)** expression was compared (n = 5). Data are representative of two independent experiments. **(I)** Heatmaps showing z-scored surface marker expression in Ly6C⁺ and Ly6C⁻ BMDEs (n = 4) stimulated with IL-5, IL-5+IFNγ, or IL-5+IFNβ. Z-scoring was performed separately for Ly6C⁺ and Ly6C⁻ subsets to highlight subset relative changes upon stimulation. Bottom annotation indicates stimulation conditions (gray = IL-5, yellow = IFNγ, green = IFNβ). **(J)** Percentage of Annexin-V+ E0771 cells after direct co-culture with IFN-stimulated Ly6C^+^ (n = 8) and Ly6C^−^ (n = 8) BMDEs. Data pooled from 2 independent experiments. (Data show individual values + median and were analysed by 2-way ANOVA for comparison of 2 or more groups. Otherwise, data show individual values + median and were analysed by Student’s t-test. Statistical significance is displayed on figures as follows: *p < 0.05, **p < 0.01, ***p<0.001, ****p<0.0001.

To validate the transcriptomic dataset, we first assessed the proposed IFN responsiveness of Ly6C^+^ TAE on protein level. FACS-sorted Ly6C^+^ and Ly6C^−^ TAEs were stimulated with IFNγ or IFNβ and PD-L1 expression was quantified relative to unstimulated counterparts (Fig. 4C). Because eosinophils express PD-L1 in an IFNγ-dependent manner^38^, PD-L1 expression served as a readout for comparison of IFN responsiveness. Consistent with the transcriptomic data (Fig. 4B), Ly6C^+^ TAEs presented with significantly higher relative PD-L1 upregulation following stimulation by either interferon, compared to their Ly6C^−^ counterparts (Supplementary Fig. 5C). Beyond PD-L1 induction, IFN stimulation broadly remodeled eosinophil activation states. In line with reports linking IFNγ to eosinophil degranulation and upregulation of antigen-presenting molecules^17^, both IFNγ and IFNβ reduced granularity and upregulated expression of MHC-I, and MHC-II (Fig. 4D). Notably, IFNγ and IFNβ promoted overlapping yet distinct eosinophil activation programs, with IFNγ favoring increase in immunomodulatory and adhesion (ICAM-1) markers, and IFNβ driving a more CD62L^+^ Ly6C^+^ phenotype (Fig. 4D).

Given the IFN-driven STAT-1 dependent Ly6C expression^37^ and the potential of both TAE subsets to overexpress Ly6C following IFN stimulation led us to investigate extent of this effect further. Intriguingly, a fraction of Ly6C^−^ tumor-associated eosinophils re-expressed Ly6C to levels comparable with native Ly6C^+^ cells, giving rise to approximately 25% and 35% of Ly6C^+^ eosinophils on average following IFNγ and IFNβ stimulation, respectively (Fig. 4E, F). To assess whether the IFN-induced Ly6C re-expression reflected functional reprogramming, IFN-stimulated eosinophils were re-gated according to their new Ly6C status into Ly6C^new−^ and Ly6C^new+^ subsets. Even though re-expression of Ly6C was not associated with the regaining of granularity or expression of MHC-I by the Ly6C^new+^ subset (Supplementary Fig. 5D, E), Ly6C^new+^ eosinophils presented with increased levels of adhesion molecules such as ICAM-1 and CD62L (Fig. 4G, H). However, the low viability of IFN stimulated TAEs (Supplementary Fig. 5F), precluded further assessment of cytotoxicity of these populations.

We therefore examined whether the BMDE model recapitulates any of the observed changes of TAEs stimulated with IFNs, since IFNs had a lesser effect on BMDE viability (Supplementary Fig. 5G). In agreement with the increased IFN responsiveness of Ly6C^+^ TAE, Ly6C^+^ BMDEs upregulated PD-L1 significantly more compared to BMDE Ly6C^−^ subset (Supplementary Fig. 5H). Of note, Ly6C^+^ BMDE induced approximately 10-times stronger relative PD-L1 expression than Ly6C+ TAE, possibly reflecting the desensitization of tumor-associated eosinophils by chronic IFN exposure within the TME. Importantly, stimulation of BMDEs with either IFNγ or IFNβ reproduced several IFN-responsive features observed in TAE (Fig. 4I), including increased expression of PD-L1, and ICAM-1 upon IFNγ stimulation and IFNβ driven upregulation of CD62L. Despite the effect on Ly6C expression was milder compared to IFN-stimulated TAE (Supplementary Fig. 5I, J), both IFNγ and IFNβ enhanced the cytotoxicity of BMDEs, with IFN stimulation restoring the cytotoxicity of Ly6C^−^ eosinophils to native Ly6C^+^ eosinophil levels (Fig. 4J).

Overall, our data demonstrate that Ly6C also serves as a marker of IFN responsiveness, with Ly6C^+^ eosinophils representing a more IFN-responsive subset. Moreover, IFNs are potent modulators of eosinophil phenotype, capable of increasing expression of activation markers and enhancing cytotoxic potential.

### IFN concentration correlates with a more active Ly6C^+^ eosinophil phenotype during ICB response

The impact of IFN on Ly6C expression and eosinophil activation ex vivo raised the question of whether IFN signaling is required to maintain the Ly6C^+^ phenotype in the TME, and/or impacts activation status in vivo. To test this, we first simultaneously neutralized IFNγ and blocked the IFNAR1 receptor which is essential for IFNα/IFNβ signaling (Fig. 5A). While blocking IFN signaling led to enlarged tumors as expected (Fig. 5B), there was no effect on overall eosinophil proportion (Supplementary Fig. 6A), or the proportion of Ly6C^+^ eosinophils (Fig. 5C). In line with the literature, IFN was essential for the expression of MHC-I and MHC-II of both Ly6C^+^ and Ly6C^−^ eosinophil subsets (Fig. 5D, Supplementary Fig. 6B), confirming that eosinophils respond to IFNs in the TME, but that loss of IFN signaling does not alter Ly6C^+^ status or expression of the canonical myeloid/activation markers in TAE (Supplementary Fig 6C). These data suggest that, whilst ex vivo eosinophils are able to respond to IFNs by upregulating receptors associated with both antigen presentation and activation (Fig. 4D, H), in the treatment-naïve breast TME these cells only require IFNs for expression of the canonical IFN-regulated molecules such as MHC-I/MHC-II.

**Figure 5.**
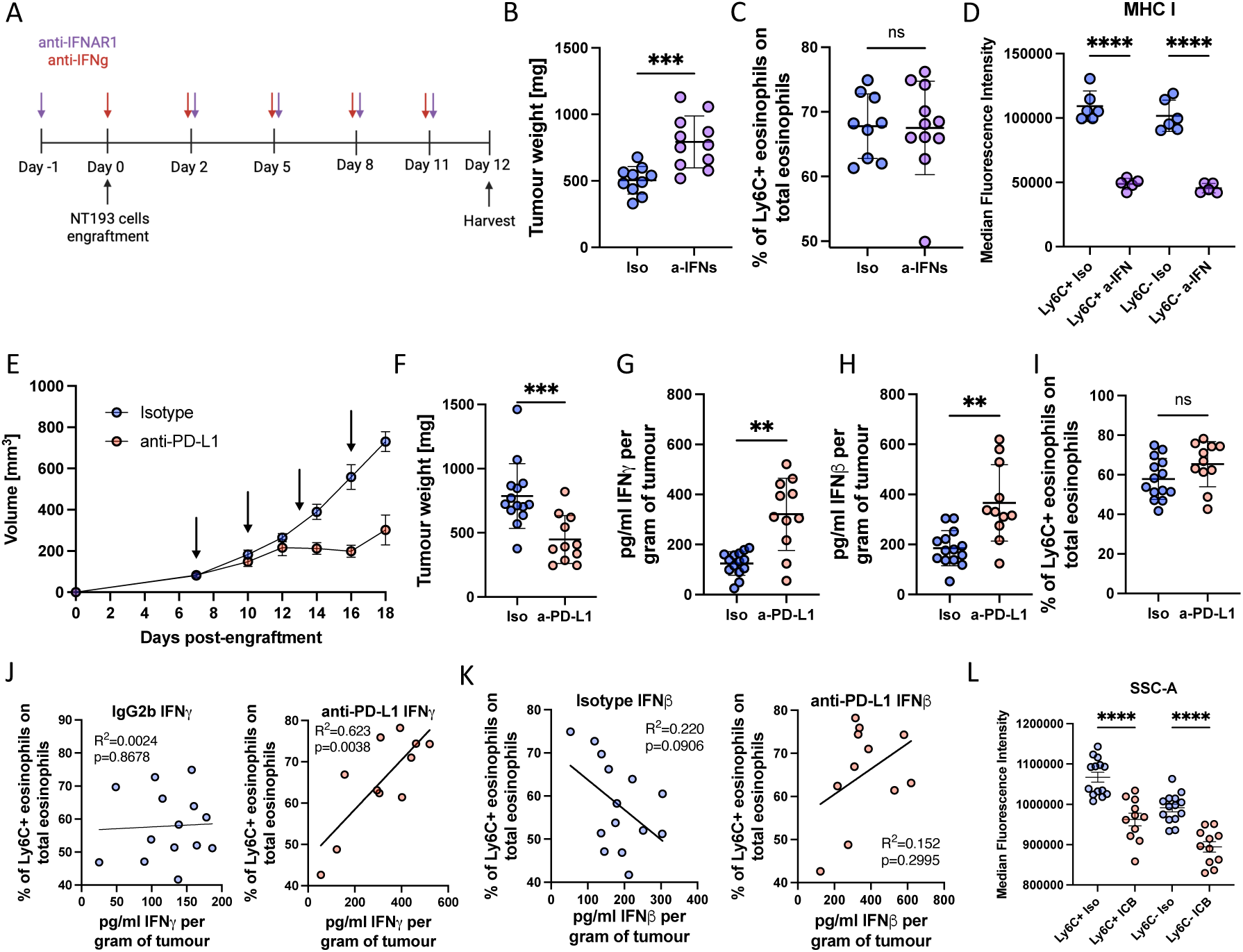
Interferons regulate the Ly6C eosinophil phenotype and cytotoxicity during anti-PD-L1 treatment. **(A)** Schematics of the experiment. NT193 tumor-bearing mice were treated with anti-IFNAR1 and anti-IFNγ antibodies (anti-IFN) or isotype control (Iso) as indicated. **(B)** Tumor weight, **(C)** proportion of Ly6C^+^ eosinophils, and **(D)** comparison of MHC-I expression on Ly6C+ and Ly6C-eosinophils (n_isotype_ = 9, n_anti-IFN_ = 11); data are representative of 2 independent experiments. **(E-L)** NT193 tumor-bearing mice were treated with anti-PD-L1 or a matched isotype control antibody. **(E)** Tumor growth with arrows indicating treatment administration, **(F)** tumor weight, **(G)** IFNγ, and **(H)** IFNβ concentration per gram of tumor. **(I)** Proportion of Ly6C+ eosinophils. **(J)** Correlation of IFNγ concentration with proportions of Ly6C+ eosinophils in isotype or anti-PD-L1 antibody-treated mice, as indicated. Data pooled from 2 experiments. **(K)** Correlation of IFNβ concentration with proportions of Ly6C+ eosinophils in isotype-treated and anti-PD-L1-treated mice, as indicated. **(L)** Comparison of SSC-A on Ly6C+ and Ly6C- eosinophils treated with isotype control or anti-PD-L1; data are pooled from 2 independent experiments. (n_Iso_=14, n_anti-PD-L1_=11). All data show individual values and mean or mean ± SD and were analysed by unpaired Student’s t-test, or by 2-way ANOVA for comparison of two and more groups. Statistical significance is displayed on figures as follows: ns > 0.05, *p < 0.05, **p < 0.01, ***p<0.001, ****p<0.0001.

Given the emerging role of eosinophils in mediating ICB responses^6,7^, the central role of IFN signaling in ICB efficacy^6^, and our findings of the IFN-responsive eosinophil subset associated with increased cytotoxicity, we next aimed to understand how eosinophils behave in an ICB-treated IFN-rich TME. NT193 tumor-bearing mice were treated with anti-PD-L1 or corresponding isotype control antibody; as expected, anti-PD-L1 ICB therapy induced tumor regression (Fig. 5E) and significantly reduced tumor burden (Fig. 5F). The concentration of both IFNγ and IFNβ was elevated in mice treated with the anti-PD-L1 treatment, confirming a successful establishment of an IFN-rich environment (Fig. 5G, H). In contrast to literature showing increased TAE in ICB treated breast tumors^6^, in this model the proportion of eosinophils in the TME was not altered following ICB treatment (Supplementary Fig. 6D). However, whilst the mean frequency of Ly6C^+^ eosinophils was unchanged in ICB treated mice (Fig. 5I), Ly6C^+^ eosinophil numbers positively correlated with IFNγ concentration in the IFN-rich TME of anti-PD-L1-treated mice, but not in the IFN-poor TME of the control cohort (Fig. 5J) and similar trends were observed with levels of IFNβ (Fig. 5K). In contrast, IFN levels did not correlate with total eosinophil levels in either of the conditions (Supplementary Fig. 6E, F). Furthermore, in an IFN-rich environment, eosinophils presented with decreased granularity, indicative of degranulated status (Fig. 5L).

These data show that IFN levels in the ICB-treated TME are significantly associated with elevated Ly6C expression and with eosinophil degranulation in vivo, in line with ex vivo data showing that the Ly6C^+^ cytotoxic TAE phenotype is driven by IFN.

## Discussion

Eosinophils are emerging as important contributors to anti-tumor immunity^39^. However, their phenotypic and functional heterogeneity within the TME remains unclear, despite recent advances in their single-cell profiling^40^. Using the eosinophil-rich NT193 breast cancer model, we identify two tumor-associated eosinophil populations distinguished by the Ly6C expression. While Ly6C^+^ eosinophils infiltrated tumors early on, Ly6C^−^ eosinophil population emerged as a hallmark of a late-stage tumor development and presented with a reduced cytotoxic potential. These findings are highly relevant, given the recent reports identifying Ly6C loss as a marker of vascular-restricted pro-metastatic neutrophil populations^41^, raising the possibility that Ly6C downregulation is a shared feature of tumor-conditioned granulocytes. In contrast to vascular-restricted neutrophils, we find that tumor-associated eosinophils are tissue resident population, with increased Ly6C expression denoting a subset of more cytotoxic and IFN-responsive eosinophil population (Figure 6). These findings help understand how different eosinophil states potentially contribute to anti-tumor immunity, including their role in mediating ICB responses^6,7,42^.

**Figure 6.**
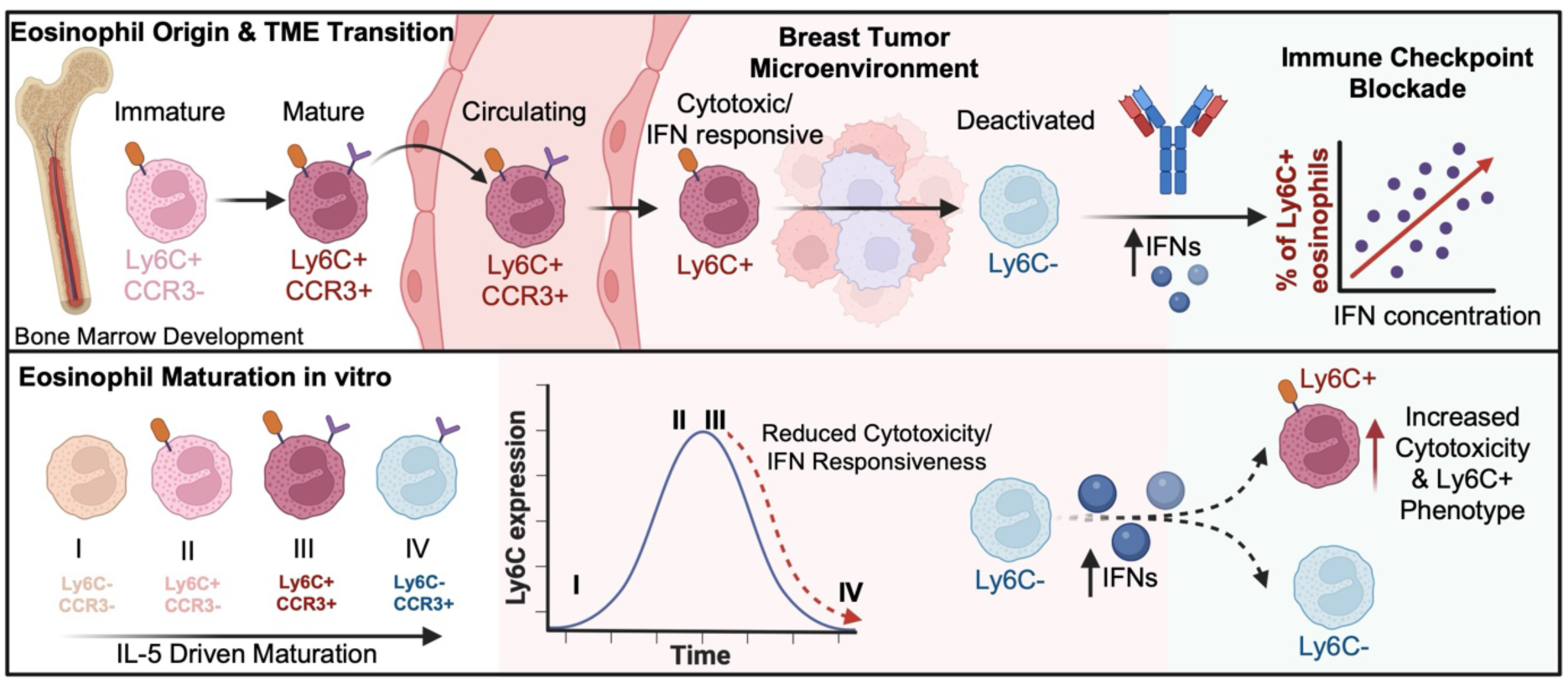
Summary of eosinophil differentiation and functional plasticity in vivo and in vitro. Eosinophils develop in the bone marrow and mature into a cytotoxic and IFN responsive Ly6C^+^ CCR3^+^ population that is present in the circulation and early-stage tumors, however loses this activation during tumor progression, resulting in a Ly6C^−^ eosinophil subset. We show that the loss of Ly6C coinciding with the loss of cytotoxicity and IFN responsiveness can be mirrored in vitro during bone marrow-derived eosinophil development. Increased levels of IFNγ and IFNβ correlate with proportions of activated Ly6C^+^ eosinophils with during immune checkpoint blockade treatment in vivo, and IFNs can enhance the activated Ly6C^+^ phenotype in vitro. Graphics created with BioRender.

We show that the well-conserved shift from a Ly6C^+^ to a Ly6C^−^ eosinophil state during tumor progression, is recapitulated in ex vivo and in vitro eosinophil differentiation assays. In line with our observations, Ly6C was recently reported to be dynamically modulated during murine eosinophilopoiesis, with intermediate developmental stages characterized by increased Ly6C expression, however mild downregulation of Ly6C towards the final developmental stage^43^. Taken together, these observations indicate that progressive Ly6C downregulation represents a shared, time-dependent feature across these distinct experimental settings, positioning Ly6C as a marker of an eosinophil-intrinsic differentiation process within environments supporting eosinophil survival.

Given the relatively low abundance of eosinophils during hemostatic conditions in vivo and the technical challenges associated with their isolation, IL-5-diven differentiation of BMDEs is commonly employed to generate sufficient eosinophil numbers for functional and mechanistic studies^36^. We found that Ly6C expression was clearly regulated during the BMDE developmental assay, where levels aligned with distinct maturation stages. By examining Ly6C together with CCR3 expression, we revealed previously unappreciated granularity of the BMDE model. Consistent with previous studies^36^, almost 100% of the culture consisted of Siglec-F+ eosinophils on day 14. However, this seemingly uniform population consists of at least four eosinophil subsets defined by distinct expression of Ly6C and CCR3, differing in granularity, maturation, cytotoxicity, and responsiveness to IFNs. This heterogeneity is particularly relevant given the widespread use of bulk BMDEs for eosinophil adoptive transfer^44–50^, cytokine stimulation^17,51^, or studying the effect of gene knockout on eosinophil development^52^. Recognizing the complexity of the BMDE culture may help with a more tailored usage of this model in the future.

BMDEs offer a robust model for studying Ly6C dynamics, although there are some phenotypic differences of these cells with TAEs, which we hypothesize stem from lack of chronic exposure to the TME. Because BMDEs mature under homeostatic conditions, IFN pathways were not differentially expressed in Ly6C^+^ BMDEs on transcriptomic level. Nevertheless, in both contexts, the loss of Ly6C is consistently associated with reduced cytotoxicity and responsiveness to IFNs. These results showing that Ly6C^−^ eosinophils lose IFN responsiveness in both TME and BMDE developmental assays, suggest that loss of IFN-responsiveness is also partially time-dependent process that is a hallmark of a more aged eosinophil state that loses cytotoxic abilities. While the mechanism behind the Ly6C transition in either of these models remains unclear, it overall highlights the importance of maintaining a pool of Ly6C^+^ eosinophils to better control tumor growth.

Several studies demonstrated that IFNγ is a potent regulator of eosinophil activation and cytotoxic potential in both human and murine assays^9,10,17,38,53–56^. Consistent with this body of work, IFNγ stimulation of both Ly6C^+^ and Ly6C^−^ TAE and BMDEs led to degranulation, upregulation of PD-L1, MHC molecules, and ICAM-1, resulting in enhancement of cytotoxicity against cancer cell lines. The importance of IFNγ-driven eosinophil activation in anti-tumor immunity is further supported by reports showing that adoptive transfer of IFNγ stimulated eosinophils reduces metastatic burden^9^. In our study, the unbiased bulkRNA-seq analysis of TAEs highlighted the strong enrichment of both IFNγ and IFNβ signatures among the Ly6C^+^ eosinophils, suggesting that activated eosinophils in the TME are also modulated by type I interferons. Despite the evidence that eosinophils are responsive to type I interferons^57^, no studies have directly examined the role of IFNβ in modulating eosinophil phenotype and function. Here we show that IFNβ can act in a complementary manner to IFNγ, with increased capacity to induce the Ly6C expression ex vivo. The ability of IFNβ to lead eosinophils into the more active Ly6C^+^ state more efficiently than IFNγ is particularly interesting given the more pronounced cytotoxic activity observed upon IFNβ stimulation. These findings raise the broader question of how type I and type II interferons jointly shape eosinophil phenotype and cytotoxicity within the complex TME.

In the context of eosinophils emerging as mediators of ICB^6^, the selective IFN-responsiveness delegated to the cytotoxic Ly6C^+^ subset, suggests their involvement in successful ICB response, which is commonly associated with IFN induction^58–60^. Here, we show that in a treatment-naïve setting, neither type I nor type II interferon signaling is detrimental for the presence of the Ly6C^+^ subset in the TME, although depletion of IFNs leads to a reduction of the MHC-I/MHC-II^+^ phenotype. Whether eosinophils lose the ability to upregulate markers associated with cytotoxic potential, and/or require additional factors to activate this response remains to be addressed. In contrast, a 2-fold increase of both IFNγ and IFNβ during ICB treatment leads to a positive correlation of the Ly6C^+^ eosinophils with IFN levels, suggestive of IFN-driven maintenance of the more active eosinophil subset during effective ICB treatment.

To conclude, our study identifies Ly6C as a key marker of eosinophil maturity and activation, with the TME leading eosinophils toward a less IFN-responsive Ly6C^−^ state (Figure 6). Our results contribute to the role of IFNs as potent eosinophil activators and suggest a new mechanism by which IFNs support ICB responses through maintenance of the Ly6C^+^ eosinophil pool.

## Methods

### Mice

FVB, Balb/cAnNCrl, and C57BL/6J wild-type mice were purchased from Charles River; all animals underwent no experimental procedures for at least 7 days post-arrival. All mice were housed under pathogen-free conditions in individually ventilated cages. A maximum of 7 mice were maintained per cage, and the mice had access to water and food ad libitum. In all experiments, female mice were orthotopically engrafted at 8–13 weeks of age. Schedule 1 method of exposure to increasing CO_2_ concentration was applied, followed by cervical dissociation at the final time point in all animal experiments. All mouse experiments were performed in accordance with the UK Home Office guidelines under project licenses PP3609558 and PA780D61A.

### Cancer cell lines

The NT193 murine breast cancer cell line, derived from a spontaneous MMTV-NeuNT primary tumor, was provided by Dr. Gertraud Orend. The 4T1 and E0771 cancer cell lines were a gift from Dr. Ilaria Malanchi. NT193 cells were cultured in Dulbecco’s Modified Eagle Medium (DMEM) (cat. 10566016, Gibco) supplemented with 10% fetal bovine serum (FBS) (cat. 10500064, Gibco), 1% Penicillin-Streptomycin (P/S) (cat. 15140122, ThermoFisher), and 10 μg/mL puromycin (cat. A1113803, Gibco). The 4T1 and E0771 cell lines were cultured in DMEM (cat. 41965039, Gibco) supplemented with 10% FBS and 1% P/S. ID8 cells were cultured in high-glucose 4.5 g/L DMEM (cat. 21969035, Life Technologies) supplemented with 4% heat-inactivated fetal bovine serum (Sigma and Life Technologies), 2mM glutamine (cat. 25030024, Life Technologies), ITS (cat. 41400045, Life Technologies) (10 µg/mL insulin, 5.5 µg/mL transferrin, and 6.7 ng/mL sodium selenite). Cells were passaged using 0.1% Trypsin-EDTA (cat. 15400054, Gibco). All cells were maintained at 37°C with 5% CO_2_ in a humidified atmosphere. All cell lines were tested for mycoplasma with MycoAlert^®^ Mycoplasma Detection Kit (cat. LT07-318, Lonza) according to the manufacturer’s instructions.

### Tumor models and administration of treatments

The NT193 cell line was used for orthotopic engraftment at full confluence, 4T1 and E0771 cell lines were used at 70-80% confluence for orthotopic engraftment. The NT193 and 4T1 cell lines were harvested with 0.25% Trypsin-EDTA, and washed three times with PBS (cat. 10010023, Gibco) prior to resuspension at the desired concentration in PBS. E0771 cells were harvested by ice-cold PBS, washed three times, and resuspended in PBS to the desired concentration. Before the orthotopic mammary fat pad injection, mice were anaesthetized with isoflurane (IsoFlo, Zoetis), the 4^th^ right mammary fat pad area was shaved and sterilized with betadine (cat. 3030440, Vidine antiseptic solution 10% w/w). To study primary breast tumor development, NT193 (5×10^6^ cells), 4T1 (5×10^4^ cells), and E0771 (1×10^5^ cells) cell lines were orthotopically grafted in 100 μL of PBS into the 4^th^ right mammary fat pad of FVB, Balb/c and C57/Bl6 female mice, respectively. Tumor-bearing mice were housed on alpha-dry bedding to avoid irritation of the skin during tumor development and tumor growth was monitored every 2-3 days, starting 4 days after the tumor induction until the final time point or up to reaching the humane endpoint (12 mm in any dimension). Tumor volume was calculated by the formula V = (length×width×width)/2. Treatment-naïve mice with spontaneously regressing tumor volume were excluded from the analysis. ID8 Pten1.14 cells were injected intraperitoneally with 5×10^6^ in 200 µl PBS. Mice were monitored regularly and killed upon reaching moderate severity limit as permitted by the Project Licence limits, which included weight loss, reduced movement, hunching, jaundice and abdominal swelling.

For all experiments involving treatments, mice were allocated to treatment groups based on their weight, age, and tumor size, when applicable, at the start of the treatment. For the eosinophil depletion, 20 μg per mouse of anti-Siglec-F antibody (cat. MAB17061-500, R&D Systems) and IgG2a isotype control (cat. BE0089, BioXCell) was intraperitoneally administered every 3 days, starting on the day of orthotopic engraftment. For the depletion of IFNγ and IFNβ signalling in vivo, combination of Ultra-LEAF Purified anti-mouse IFNAR-1 (cat. 400198, BioLegend) and Ultra-LEA Purified anti-mouse IFN-γ (cat. 505848, BioLegend) or a control Ultra-LEAF Purified Mouse IgG1, Isotype antibody (cat. 400198, BioLegend) was administered by intraperitoneal injection; 1.5 mg of anti-IFNAR-1 antibody day prior to the tumor induction, 250 μg of anti-IFNγ on the day of orthotopic engraftment, and 300 μg of anti-IFNAR1 and 250 μg of anti-IFNγ, every 3 days from the day 2 post-engraftment onwards. For the anti-PD-L1 treatment, Ultra-LEAF Purified anti-mouse CD274 (cat. 124339, BioLegend) or Ultra-LEAF Purified Rat IgG2b antibodies (cat. 400672, BioLegend) were administered every 3 days by intraperitoneal injection at a final dose of 10 mg/kg, starting on day 7 post-engraftment.

### In vivo CD45-APC labelling

NT193 tumor-bearing mice were intravenously injected with 5 μg of anti-CD45 (clone 30-F11, BioLegend) and culled by cervical dislocation 2 minutes after. Tumor, blood, and bone marrow were collected and processed for flow cytometry as described below.

### Tissue processing

At experimental timepoint or when tumors reached 12 mm in any dimension, tumors were collected into ice-cold PBS and manually dissociated using scissors, resuspended in tumor digestion media (Roswell Park Memorial Institute 1640 medium (RPMI), 10% FBS, 500 μg/mL Liberase TM (cat. 5401127001, Roche), 100 μg/mL DNAse I (cat. 11284932001, Sigma)) and digested for 30 minutes at 37°C under gentle agitation. Following enzymatic digestion, tumor suspensions were mechanically passed through 70 μm cell strainers (cat. 83.3945.070, Sarstedt) and washed once with FACS buffer (PBS supplemented with 5% FBS, 10 μg/mL DNAse I) to obtain a single cell suspension. Identical protocol was used for processing single of healthy mammary fat pads. For time course experiments analysing blood samples in eosinophil depletion experiment, blood was collected by tail vain puncture using Jaytec Glass Micro-Haematocrit Tubes (cat. 12306297, Fisher Scientific) and resuspended in 25µl of 0.5 M EDTA (cat. 15575020, ThermoFisher) straight after. For experiments comparing matched tumor and blood samples in Ly6C^−^ eosinophil tracking experiment, blood was collected by cardiac puncture into an EDTA-coated tube (cat. 367527, BD). All samples were treated with ACK buffer for 10 minutes at room temperature, and lysis was stopped by ice-cold PBS. Bone marrow used for flow cytometry analysis was collected by flushing the femur bones with 5 mL of cRPMI. All cell suspensions were spun down at 1500 rpm for 5 minutes, resuspended in FACS buffer and kept at 4°C for flow cytometry staining.

### Flow cytometry and sorting

All samples stained for flow cytometry analysis were plated into V-shaped 96-well plates at a maximum 2×10^6^ cells per well for tissue samples, a minimum of 5×10^4^ cells per well was plated for eosinophil developmental assays and IFN stimulations, and a minimum of 1.5×10^4^ cells per condition was plated for assays with FACS-sorted eosinophils that underwent development ex vivo. All cells were incubated with TruStain FcX (cat. 101319, ThermoFisher) at 1:200 dilution for 15 min at 4°C. Blocked samples were stained with fixable viability dyes (LIVE/DEAD Fixable Blue, Near-IR, or Yellow Dead Cell Stain Kits; cat. L23105, L34975, L34959, ThermoFisher) in PBS for 30 minutes at 4°C, according to manufacturers’ instructions. For surface staining, cells were incubated with antibodies against extracellular antigens for 30 minutes at 4°C. Cells were either acquired fresh in FACS buffer, or were fixed with Fixation buffer (cat. 420801, BioLegend) for 15 minutes, washed and stored in FACS buffer at 4°C for a maximum of 4 days.

For fluorescence-activated cell sorting (FACS) experiments, single cell suspensions after digest were enriched with mouse anti-Siglec-F (cat. 130-118-513, Miltenyi magnetic micro-beads using MS columns (cat. 130-042-201, Miltenyi) in combination with OctoMACS™ Separator (Miltenyi) following manufacturer’s instructions. Enriched cells were further stained with LIVE/DEAD Fixable Near-IR Dead Cell Stain Kit and surface receptors as described above, and resuspended in PBS, 2 mM EDTA, and 0.5% FBS before sorting with a 100μm nozzle size on Aria III.

All data were acquired by LSRFortessa X-20 (BD Biosciences) or Aurora (Cytek) spectral flow cytometer using DIVA or SpectroFlo software, respectively. Flow cytometry data was analysed using OMIQ (Dotmatics) and FlowJo v10.9.0.

### ELISA assays

Tumor supernatants were collected following the mechanical dissociation into 1 mL of DMEM media, prior to the enzymatic digestion at 37°C. Samples were spun down for 10 minutes at 500xG, and the supernatant was stored at −20°C. IFNγ and IFNβ concentrations were quantified by sandwich BioLegend’s ELISA MAX Deluxe Set (cat. 430804, cat. 439404, BioLegend) following the manufacturer’s protocol. Concentrations were normalised to tumor weight according to the standard curve.

### Bone marrow-derived eosinophil development

To obtain eosinophils from primary murine bone marrow cells, cells were cultured as previously described^36^. Freshly isolated bone marrow cells were cultured in RPMI 1640 supplemented with 20% FBS, 25 mM HEPES, 1% P/S, 2 mM glutamine (cat. 25030-024, Gibco), 1x NEAA (cat. 11140-035, Gibco), and 1 mM sodium pyruvate (cat. 11360070, Gibco) (eosinophil media) for 4 days supplemented with 100 ng/mL mSCF (cat. 250-03, PeproTech) and 100 ng/mL mFLT3-Ligand (cat. 250-31L, PeproTech), followed by 10 ng/mL IL-5 (cat. 215-15, PeproTech) from day 4 onwards. Half of the media was replaced every 2 days; during all media replacements, eosinophils were spun at 300xG for 7 minutes. On days 4 and 8, cells were plated into a new flask to minimise the culture of adherent cells. Cell culture was kept at 1×10^6^ cells/mL confluence. In experiments with bone marrow-derived eosinophils, both Bl6 male and female mice between 8-13 weeks were used.

### Eosinophil ex vivo cultures

Eosinophils from NT193 tumors or bone marrow cultures were FACS-sorted into eosinophil media supplemented with IL-5, washed once with RPMI, and kept in culture for the indicated number of days in presence of 10 ng/mL IL-5. For IFN stimulation experiments, sorted eosinophils were cultured in presence of 10 ng/mL IL-5, supplemented with either IFNγ (10 ng/mL) (cat. 575304, BioLegend), IFNβ (10 ng/mL) (cat. 581304, BioLegend), or in absence of IFNs for 24 hours and analysed by flow cytometry as described above.

### Eosinophil cytotoxic assays

For eosinophil cytotoxic assays with unstimulated tumor sorted Ly6C^+^ and Ly6C^−^ eosinophils, eosinophils were in a direct co-culture with the NT193 cell line at a 1:1 ratio for 3 days. For eosinophil cytotoxic assays with unstimulated bone marrow-derived eosinophils, sorted CCR3^+^ Ly6C^+^ and CCR3^+^ Ly6C^−^ eosinophils were in a direct co-culture with the E0771 cell line at a 1:1 ratio for 1 day. In assays using bone marrow-derived eosinophils stimulated with IFNs, eosinophils were stimulated as described above, washed once with RPMI, and were in a direct co-culture with the E0771 cell line at a 1:1 ratio for 1 day. All cytotoxic assays were performed in eosinophil media supplemented with 10 ng/mL of IL-5.

Cytotoxic assays were stopped by the collection of cells into a separate 96-well V-shaped plate, cancer cells were washed once with PBS and collected with 0.25% Trypsin-EDTA. During all washing steps, supernatants were collected and pooled. Cells were blocked with 1:200 Fc-block, stained with fixable viability dye and anti-Siglec-F or anti-CD45 antibody as described above. All following steps were performed in 1X Annexin-V binding buffer (cat. 422201, BioLegend) per manufacturer’s instructions. Samples were washed once, stained with Annexin-V (cat. 640920, BioLegend) at 1:60 for 15 minutes at room temperature in the dark, washed, and acquired immediately after the final wash. SiglecF^−^/CD45^−^ cells were considered as cancer cells and were further analysed for Annexin-V and Live/dead staining, with double positive cells representing the apoptotic population.

### Imaging

Harvested murine tumors were fixed in PFA-based fixative Antigenfix (cat. P0014, DiaPath) for 12-24 hours at 4°C. Tumors were afterwards washed twice with PBS for 1 hour at 4°C and dehydrated in 30% sucrose (cat. S9378-5KG, Sigma) for 24-48 hours. Following dehydration, tumors were embedded in Tissue-Tek OCT compound (cat. 16-004004, Tissue-Tek), and frozen using methanol with dry ice. OCT blocks were sectioned using Leica CM3050 S Cryostat, sections of 5-10 μm were mounted on positively charged VWR SuperFrost Plus, Adhesion Slides (cat. 631-0108, Avantor) and stored at −80°C. For immunofluorescent staining, sections were incubated for 5 minutes at room temperature, rehydrated with PBS and blocked with PBS with 2% FBS, 0.1% Triton X-100 (cat. X100, Sigma), 5% donkey serum (cat. D9663, Sigma), and TruStain FcX 1:200 (blocking solution) for 4 hours in a humid chamber at room temperature. Blocked samples were incubated with primary antibodies overnight at 4°C. Samples were washed three times with blocking buffer and incubated with secondary antibodies diluted in blocking solution at room temperature for 4 hours in a humid chamber. In experiments with nuclear staining, sections were stained with 1 μg/mL DAPI (cat. D3571, ThermoFisher) for 15 minutes at room temperature and washed three times with PBS. Samples were mounted using FluorSave reagent (cat. 345789, Millipore) and imaged on ZEISS Axioscan 7 slide scanner unless stated otherwise.

For Cytospin imaging, up to 100000 cells in 100 μL were cytospun with Shandon Cytospin 3 Cytocentrifuge at 300xG for 3 minutes, with Scientific CytoSep Filter Papers for Shandon Cytospin Centrifuges (cat. 22-045-305, ThermoFisher) onto a positively charged slide (cat. 631-0108, VWR). Slides were left to air dry, fixed with ice-cold fixation buffer for 15 minutes at room temperature, washed, and left to air dry overnight. Haematoxylin and eosin staining was performed by an autostainer Sakura Tissue Tek DRS. Briefly, slides were fixed in methanol (cat. 322415, Sigma) for 5 seconds, air dried, stained with Harris Haematoxylin (cat. 3801560E, Leica) for 11 minutes, washed, incubated with acid alcohol (0.1% hydrochloric acid (cat. 10763124, Fisher Scientific), 70% absolute ethanol in distilled water) for 40 seconds to remove excess Haematoxylin, washed, and incubated with ammoniated water (0.3% ammonia solution (cat. 87766.290, VWR) in tap water) to enhance contrast staining. Slides were then washed in tap water and stained with eosin (cat. RBC-0100.00A, CellPath) for 2.5 minutes, prior to three washes with tap water and dehydration in 100% ethanol and xylene. Dehydrated slides were cover-slipped using an automatic Sakura Tissue-Tek Glas Automated Glass Coverslipper and imaged on ZEISS Axioscan 7 slide scanner. All images were analysed using QuPath v0.5.0.

### Bulk RNA sequencing

Tumor-associated eosinophils from NT193 tumors (20000 cells per condition – Ly6C^+^ and Ly6C^−^) and bone marrow-derived eosinophils of Bl6 mice (100000 cells) were FACS sorted as described above and resuspended in RLT (cat. 79216, Qiagen) with 1% β-mercaptoethanol (cat. M6250, Sigma). Samples were snap frozen on dry ice and RNA extraction, library preparation, sequencing, quality control and read processing were performed by Azenta Life Sciences using an ultra-low input RNA-seq workflow for tumor-associated eosinophils and a standard bulk RNA-seq pipeline for bone marrow-derived eosinophils. Sequencing was conducted on the Illumina (2×150 bp, ∼20 million paired-end reads per sample). Reads were trimmed with *Trimmomatic* (v0.36) and quality-checked with FastQC. Alignment to the Mus musculus reference genome (GRCm38, available on ENSEMBL) was performed using *STAR aligner* (v2.5.2b). Gene-level read counts were generated with *featureCounts* (Subread v1.5.2) using exon-overlapping, uniquely mapped reads.

All subsequent analyses were conducted in R version 4.4.0 within RStudio. Gene expression analysis was performed on raw counts. Lowly expressed genes were filtered out by removing genes with a total count of ≤400 across all samples. Ensembl gene identifiers were mapped to mouse gene symbols using the *org.Mm.eg.db* package (v3.19.1) via the *AnnotationDbi* package (v1.66.0). Genes lacking annotations were excluded. Differential gene expression analysis was performed using *DESeq2* (v1.44.0), with tumor identity included as a covariate to account for paired samples, with the primary comparison being between Ly6C^+^ and Ly6C^−^eosinophils. Log2 fold changes and Benjamini–Hochberg adjusted p-values (padj) were calculated using the Wald test. Genes with padj < 0.05 were considered significantly differentially expressed.

Variance-stabilising transformation (VST) was applied to the DESeq2 object for dimensionality reduction and visualisation. Principal component analysis (PCA) was performed using *plotPCA* from DESeq2 and visualised with *ggplot2* (v3.5.2). Volcano plots were generated with the *EnhancedVolcano* package (v1.22.0). For visualisation, the top 20 differentially expressed genes (DEGs) were selected based on descending padj values. Transformed expression values were z-score scaled across genes, and heatmaps were plotted using *pheatmap* (v1.0.13). Fold changes were also calculated per tumor to compare Ly6C^+^ versus Ly6C^−^eosinophils within individual tumors. Gene set enrichment analysis (GSEA) was conducted using the *clusterProfiler* package (v4.12.6). Genes were ranked by log₂ fold change and mapped to Entrez IDs. The *gseGO* function was used to identify enriched Gene Ontology (GO) terms across biological process, molecular function, and cellular component categories altogether, with significance defined as padj < 0.05. For bone-marrow–derived eosinophil (BMDE) samples, GSEA was conducted using *gseGO* restricted to the GO biological process (BP) ontology. Redundant GO terms were reduced using *simplify* (also from clusterProfiler), producing a non-redundant set of enriched pathways prior to extracting core enrichment genes and gene counts. For selected GO terms of interest, core enrichment genes were extracted and annotated. Heatmaps of these gene sets were generated using z-score-scaled expression values.

## Data analysis

Unless otherwise specified, statistical analysis was performed using GraphPad Prism v10.5. One-way or two-way ANOVA using Holm-Šídák correction, or unpaired Student’s t-test were applied as indicated in figure legends. Statistical significance was set at p-value < 0.05.

## Acknowledgments

We would like to thank Azenta Life Sciences for the generation of the sequencing data, Jonathan Webber for the assistance with cell sorting, Albertino Bonifacio and Karolina Kaczkowska for help with in vivo experiments, Helen Byrne, Joe Pitt-Francis, Philip Maini, and Vedang Narain from the Mathematical Institute, University of Oxford for their contribution to the funding acquisition.

## Author contributions

**Z.V.** - Conceptualization, Methodology, Investigation, Formal analysis, Visualization, Funding acquisition, Writing - original draft, Writing – review & editing; **M.P.** - Conceptualization, Methodology, Writing – review & editing; **G.J.M.** - Investigation, Writing – review & editing; **L.K.J.** - Investigation, Formal analysis, Writing – review & editing; **V.C.** - Investigation, Writing – review & editing; **S.S.** - Investigation, Formal analysis, Writing – review & editing; **I.A.M.** - Supervision, Writing – review & editing; **A.S.** - Investigation, Writing – review & editing; **A.L.H.** - Conceptualization, Funding acquisition, Project administration, Supervision, Writing – review & editing; **A.G.** - Conceptualization, Methodology, Supervision, Writing – original draft, Writing – review & editing; **K.S.M.** - Conceptualization, Funding acquisition, Project administration, Supervision, Writing – original draft, Writing – review & editing

## Data and materials availability

All data associated with this study are present in the paper or the Supplementary Materials. Bulk RNA-seq data of bone marrow-derived eosinophils and tumor-associated eosinophils will be made available online. Additional materials will be provided upon reasonable request to the corresponding author.

## Supplementary Figures

**Supplementary Figure 1.**
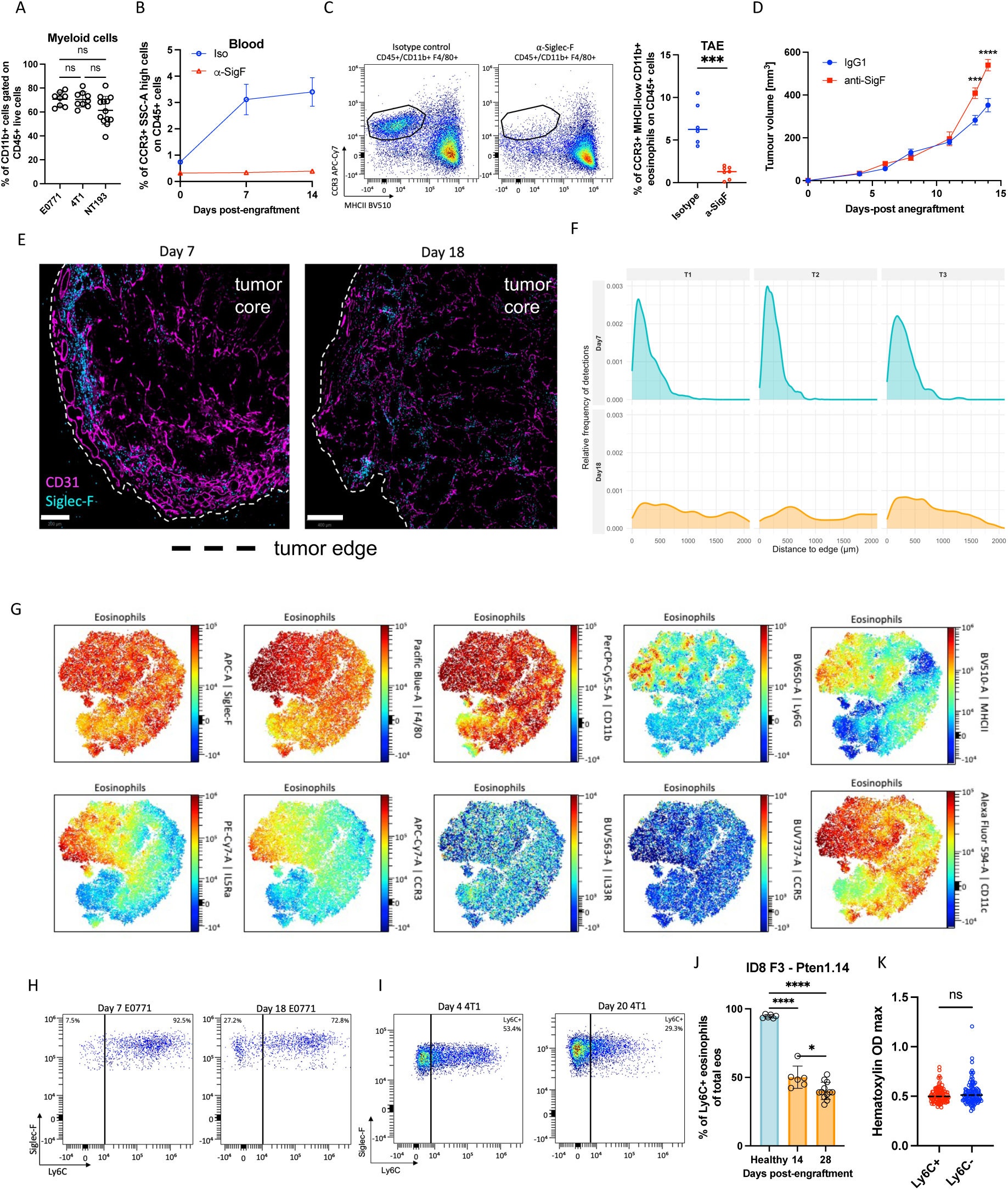
The tumor microenvironment shapes eosinophils phenotype. **(A)** Infiltration of CD11b+ myeloid cells at the final time point in the NT193 (day 18, n=11), 4T1 (day 17-20, n=9), and E0771 (day 18-22, n=8) orthotopic mammary tumors engrafted into the 4^th^ mammary fat pad of FVB, Balb/c, and C57/BL6 females, respectively. Data pooled from 2 independent experiments. **(B-D)** The NT193 tumor-bearing mice that were treated with an isotype control (n = 16) or an anti-Siglec-F antibody(n=15). **(B)** Percentage of eosinophils (Ly6G^−^ CCR3^+^, SSC^high^) of all immune cells in the blood analyzed by flow cytometry. Time course representative of one experiment, final blood confirmation representative of 3 independent experiments. **(C)** Representative flow cytometry plot of myeloid cells showing tumor-associated eosinophil gating strategy based on CCR3 and MHC-II expression and quantification of tumor-associated eosinophils (CCR3+ MHC-II-low) analyzed by flow cytometry at the final time point. **(D)** Tumor growth curve of isotype and anti-Siglec-F treated mice. Data representative of 2 independent experiments. **(E)** NT193 tumors grafted in wild-type female mice for the indicated number of days were OCT frozen and stained with Siglec-F and CD31. Scale bar day 7 = 200 µm, scale bar day 18 = 400 µm, dashed line represents tumor edge; data representative of 3 independent experiments. **(F)** Quantification of eosinophil distance to tumor edge in NT193 tumors on day 7 and day 18. Data representative of 3 independent experiments, with at least 3 tumors per condition within each experiment. **(G)** NT193 tumor-associated eosinophils were analyzed by spectral flow cytometry on day 18. Eosinophils were analyzed by opt-SNE analysis, with heatmaps of individual marker expression imposed on the opt-SNE map of the eosinophil population. **(H-I)** Representative flow cytometry plots of Ly6C expression of the tumor-infiltrating eosinophils (Siglec-F+, Ly6G-) from E0771 **(H)** and 4T1 **(I)** early and late tumors, as indicated. **(J)** Proportion of Ly6C+ eosinophils of the total eosinophil population in healthy omentum and ID8 omental tumours induced by intraperitoneal injection of ID8 Pten1.14 cells and harvested on indicated days, analyzed by flow cytometry. **(K)** Quantification of hematoxylin optical density of cytospun eosinophils (n_Ly6C+_ = 106, n_Ly6C-_ = 107). Quantification representative of 5 out of 6 tumors. All data show individual values and mean or mean ± SD and were analysed by unpaired Student’s t-test or by 2-way ANOVA for comparison of two and more groups. Statistical significance is displayed on figures as follows: ns > 0.05, *p < 0.05, **p < 0.01, ***p<0.001, ****p<0.0001.

**Supplementary Figure 2.**
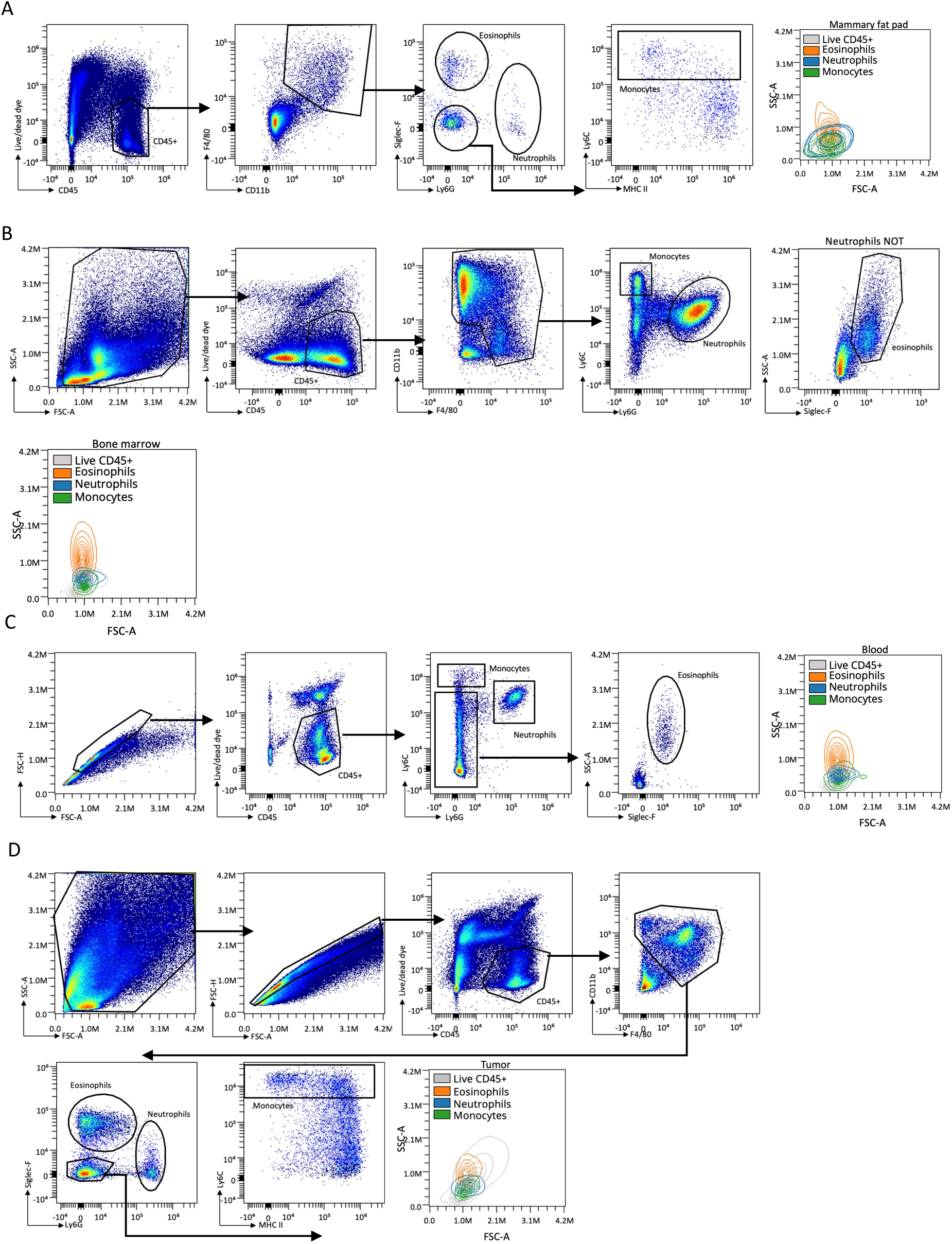
Gating strategies used to analyze flow cytometry data. **(A)** mammary fat pad, **(B)** bone marrow, **(C)** blood, **(D)** representative of tumors on both day 7 and day 18. Associated overlayed counterplots serve for the comparison of physical properties of the gated myeloid population.

**Supplementary Figure 3.**
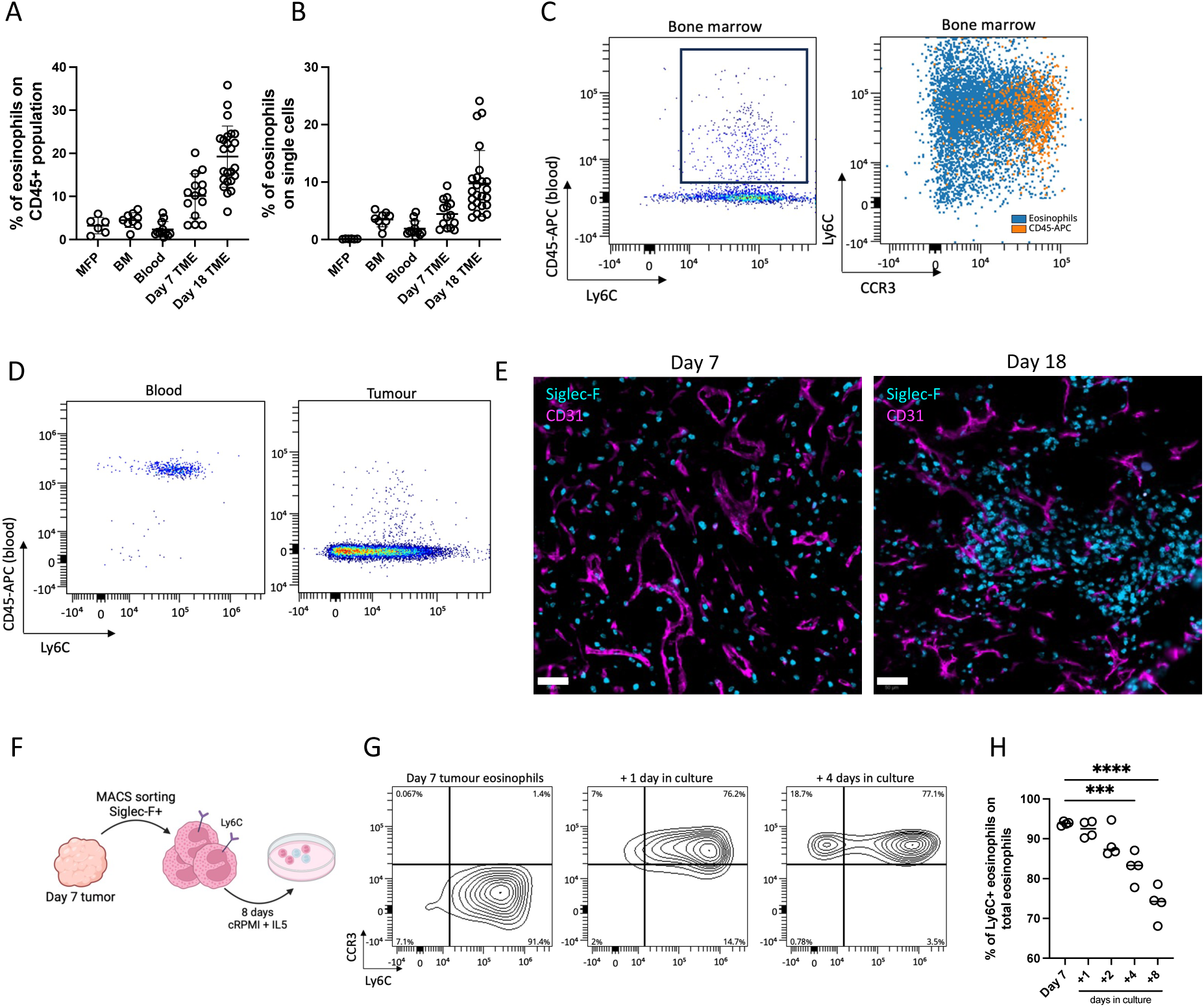
Eosinophils represent a resident population in NT193 tumors. **(A, B)** Proportion of eosinophils to all CD45^+^ immune cells **(A)** and all single cells **(B)** of healthy mammary fat pad (MFP, n = 6), bone marrow of mice bearing tumors for 18 days (BM, n = 9), blood of mice bearing tumors for 18 days (n = 7), NT193 tumors 7 days post-engraftment (Day7 TME, n = 14), and NT193 tumors 18 days post-engraftment (Day18 TME, n = 24). **(C)** Representative density plot of CD45-APC labelled bone marrow eosinophils (orange) of tumor-bearing mice compared to resident bone marrow eosinophils not labelled with the anti-CD45-APC antibody (blue). **(E)** Representative flow cytometry plot of CD45-APC labelling efficiency of blood and tumor eosinophils of NT193 bearing mice, as indicated. **(E)** Immunofluorescent staining of Siglec-F+ eosinophils (cyan) and CD31+ vasculature (magenta) of NT193 tumours on the indicated days. Scale bar = 50 µm. **(F)** Schematics of experimental design, eosinophils were MACS isolated on day 7 and cultured in the presence of IL-5. Graphics created with BioRender. **(G)** Representative flow cytometry contour plots of Ly6C and CCR3 expression of MACS isolated eosinophils from NT193 tumors cultured for the indicated number of days. **(H)** Percentage of Ly6C+ eosinophils in culture. Data representative of 2 independent experiments. All data show individual values and mean or mean ± SD and were analysed by one-way ANOVA. Statistical significance is displayed on figures as follows: ns > 0.05, *p < 0.05, **p < 0.01, ***p<0.001, ****p<0.0001.

**Supplementary Figure 4.**
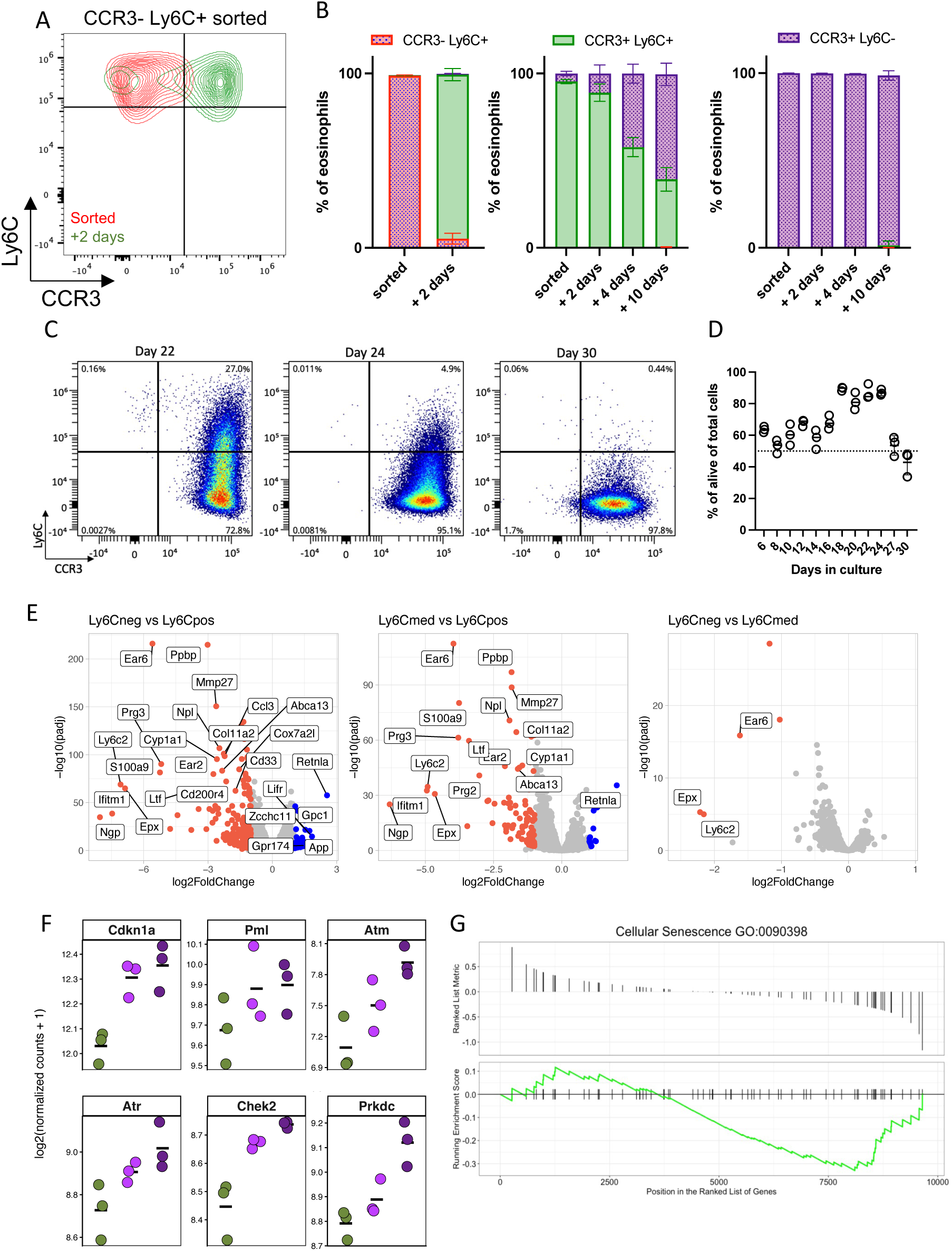
Bone marrow-derived eosinophils naturally transition to CCR3+ Ly6C-state. **(A)** Eosinophils were derived from bone marrow progenitors of C57/Bl6 mice. Representative contour plot of CCR3 and Ly6C expression on BMDE CCR3- Ly6C+ eosinophils after sorting (red) and after 2 days in culture (green). **(B)** Quantification of eosinophil development from CCR3- Ly6C+, CCR3+ Ly6C+ and CCR3+ Ly6C- cells as indicated. Data representative of 2 independent experiments**. (C)** Representative flow cytometry plots of CCR3 and Ly6C expression of BDE cultured in the presence of IL-5 up to day 30. **(D)** Proportion of Siglec-F+ bone marrow-derived eosinophils of the total viable population on indicated days. **(E-G)** FACS-sorted BMDEs analyzed by bulk-RNA sequencing. **(E)** Volcano plot analyses of Ly6C^+^, Ly6C^med^, Ly6C^−^ BM-derived eosinophils analyzed by bulkRNA-seq, as indicated. **(F)** Log2 normalized gene expression of selected senescence markers. All data show individual values and mean. **(G)** Gene set enrichment analysis (GSEA) plot for the Gene Ontology biological process *cellular senescence* (GO:0090398).

**Supplementary Figure 5.**
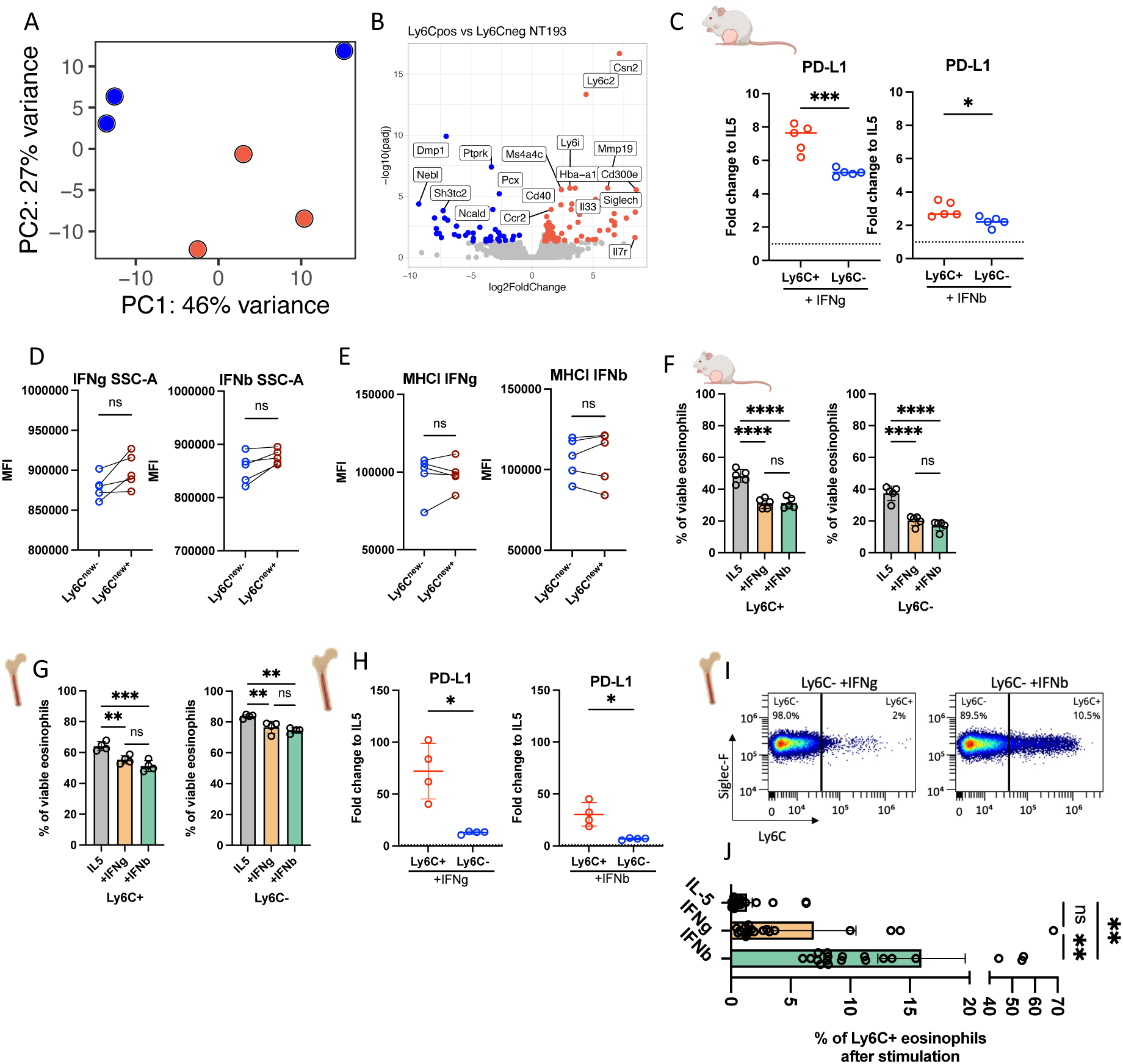
IFN stimulation affects tumor-associated and bone marrow-derived eosinophils. **(A-B)** The Ly6C^+^ and Ly6C^−^ eosinophils were FACS-sorted from the NT193 tumors and analyzed by bulkRNA-seq (n = 3). **(A)** Visualisation of principal components (PC) 1 and 2 of PCA analysis, and **(B)** volcano plot of differentially expressed genes between Ly6C^+^ (red) and Ly6C^−^ (blue) eosinophils. **(C)** Fold change of PD-L1 median fluorescence intensity of tumor-associated eosinophils cultured in the presence of IFNγ or IFNβ compared to matched IL-5 cultured eosinophils, as indicated (n_NT193_ = 5). Representative of 3 independent experiments. **(D, E)** Ly6C^−^ eosinophils stimulated with IFNγ or IFNβ were re-gated as indicated in Fig. 4E into Ly6C^new−^ and Ly6C^new+^ subsets, and granularity SSC-A **(D)**, and MHC-I **(E)** expression was compared (n = 5). Data are representative of two independent experiments. **(F, G)** Comparison of viability on **(F)** tumour-associated and **(G)** BMDE eosinophils following the IFNγ or IFNβ stimulation. (n_NT193_ = 5, n_BMDE_ = 4), data representative of at least 2 independent experiments. **(H)** Fold change of PD-L1 median fluorescence intensity of bone marrow-derived eosinophils cultured in the presence of IFNγ or IFNβ compared to matched IL-5 cultured eosinophils, as indicated (n_BMDE_ = 4). Representative of 3 independent experiments. **(I)** Representative flow cytometry plot of Ly6C expression of Ly6C^−^ BMDE stimulated with IFNγ or IFNβ, as indicated. **(J)** Proportions of Ly6C^new+^ subset of all viable BMDEs after stimulation. Data pooled from 5 independent experiments (n = 19). Data show individual values and mean and were analysed by 2-way ANOVA for comparison of 2 or more groups. Otherwise, data show individual values + median and were analysed by Student’s t-test. Statistical significance is displayed on figures as follows: *p < 0.05, **p < 0.01, ***p<0.001, ****p<0.0001.

**Supplementary Figure 6.**
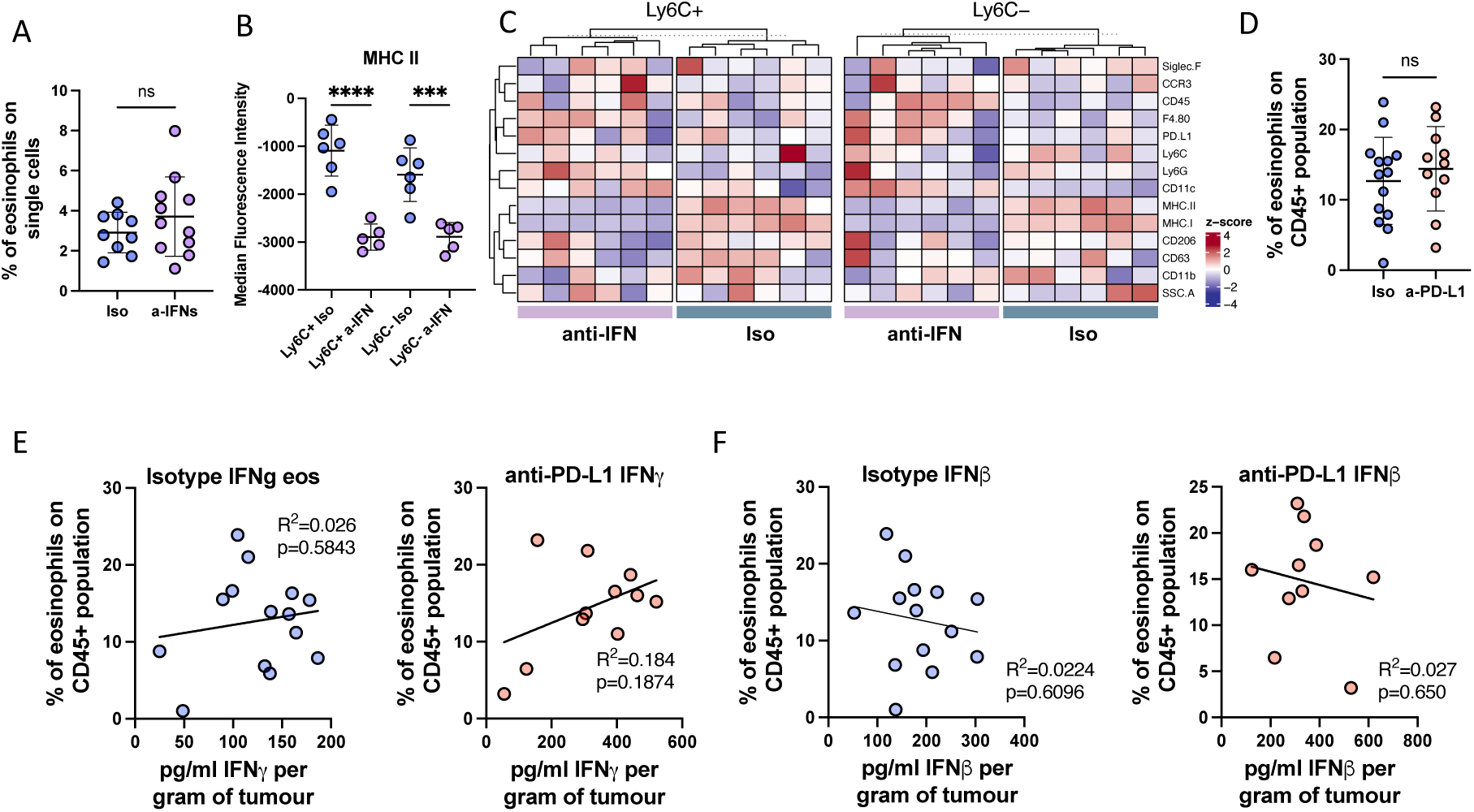
IFNs do not impact general proportions of eosinophils in the TME. **(A-C)** NT193 tumor-bearing mice were treated with anti-IFNAR1 and anti-IFNγ antibodies (anti-IFN) or isotype control (Iso). **(A)** Proportion of eosinophils on total population of viable single cells (n_isotype_ = 9, n_anti-IFN_ = 11), data pooled from 2 independent experiments, **(B)** comparison of MHC-II expression on Ly6C+ and Ly6C- eosinophils; data are representative of 2 independent experiments. **(C)** Heatmaps showing z-scored surface marker expression on Ly6C⁺ and Ly6C⁻ tumor-associated eosinophils following in vivo blockade of type I and type II IFN signaling (anti-IFN) or isotype control (Iso). Z-scoring was performed within each subset (Ly6C⁺ or Ly6C⁻) to highlight relative changes induced by IFN blockade. **(D-F)** NT193 tumor-bearing mice were treated with anti-PD-L1 or a matched isotype control antibody. **(D)** Proportion of eosinophils, **(E)** Correlation of IFNγ with proportions of eosinophils in isotype or anti-PD-L1 antibody-treated mice, as indicated, **(F)** Correlation of IFNβ concentration with proportions of eosinophils in isotype-treated and anti-PD-L1-treated mice, as indicated. Data pooled from 2 experiments. Data show individual values and mean or mean ± SD and were analysed by unpaired Student’s t-test and 2-way ANOVA for comparison of 2 or more groups. Statistical significance is displayed on figures as follows: *p < 0.05, **p < 0.01, ***p<0.001, ****p<0.0001, ns > 0.05.

